# Mechanism of REST/NRSF Regulation of Clustered Protocadherin α Genes

**DOI:** 10.1101/2021.03.06.434230

**Authors:** Yuanxiao Tang, Zhilian Jia, Honglin Xu, Lin-Tai Da, Qiang Wu

**Affiliations:** Center for Comparative Biomedicine, MOE Key Laboratory of Systems Biomedicine, State Key Laboratory of Oncogenes and Related Genes, School of Life Sciences and Biotechnology, Institute of Systems Biomedicine, Shanghai Jiao Tong University, Shanghai 200240, China

## Abstract

Repressor element-1 silencing transcription factor (REST) or neuron-restrictive silencer factor (NRSF) is a zinc-finger (ZF) containing transcriptional repressor that recognizes thousands of neuron-restrictive silencer elements (NRSEs) in mammalian genomes. How REST/NRSF regulates gene expression remains incompletely understood. Here, we investigate the binding pattern and regulation mechanism of REST/NRSF in the clustered protocadherin (*PCDH*) genes. We find that REST/NRSF directionally forms base-specific interactions with NRSEs via tandem ZFs in an anti-parallel manner but with striking conformational changes. In addition, REST/NRSF recruitment to the *HS5-1* enhancer leads to the decrease of long-range enhancer-promoter interactions and downregulation of the clustered *PCDHα* genes. Thus, REST/NRSF represses *PCDHα* gene expression through directional binding to a repertoire of NRSEs within the distal enhancer and variable target genes.

## INTRODUCTION

During early neurogenesis, the orderly acquisition and maintenance of neural identities are controlled epigenetically by de-repression of neural genes through downregulating transcriptional repressors and corepressors (1). REST (repressor element-1 silencing transcription factor), also known as NRSF (neuron-restrictive silencer factor), is a crucial repressor for neural genes (2, 3), reviewed in (4). Specifically, REST/NRSF represses the expression of numerous neural-specific genes in neural progenitors as well as non-neural tissues (5–8). In differentiated non-neural cells, REST/NRSF represses neural genes in collaboration with its corepressors (6,8–11). In embryonic stem cells, REST/NRSF is highly expressed (8, 12). During transition to neural progenitor cells (NPCs) and finally to mature neurons, REST/NRSF is degraded to minimal levels in NPCs and to an undetectable level in mature neurons (8, 13). Recent studies revealed that REST/NRSF also has a protective role in genome stability (14).

REST/NRSF contains a central DNA-binding domain with eight tandem C2H2 ZFs and two repressor domains residing in the amino and carboxyl termini, respectively (Figure 1A) (2,3,15). REST/NRSF has been shown to bind to thousands of NRSEs which can be divided into three groups: canonical, noncanonical, and half-site only motifs (16, 17). Intriguingly, canonical and noncanonical NRSEs contain very different gap sizes between the left- and right-half sites. ZF domains are small DNA-recognition units that are usually organized in tandem and there are more than 800 ZF transcription factors in the human genome (18–21). The DNA-recognition mechanisms of ZF proteins are largely unknown.

**Figure 1.**
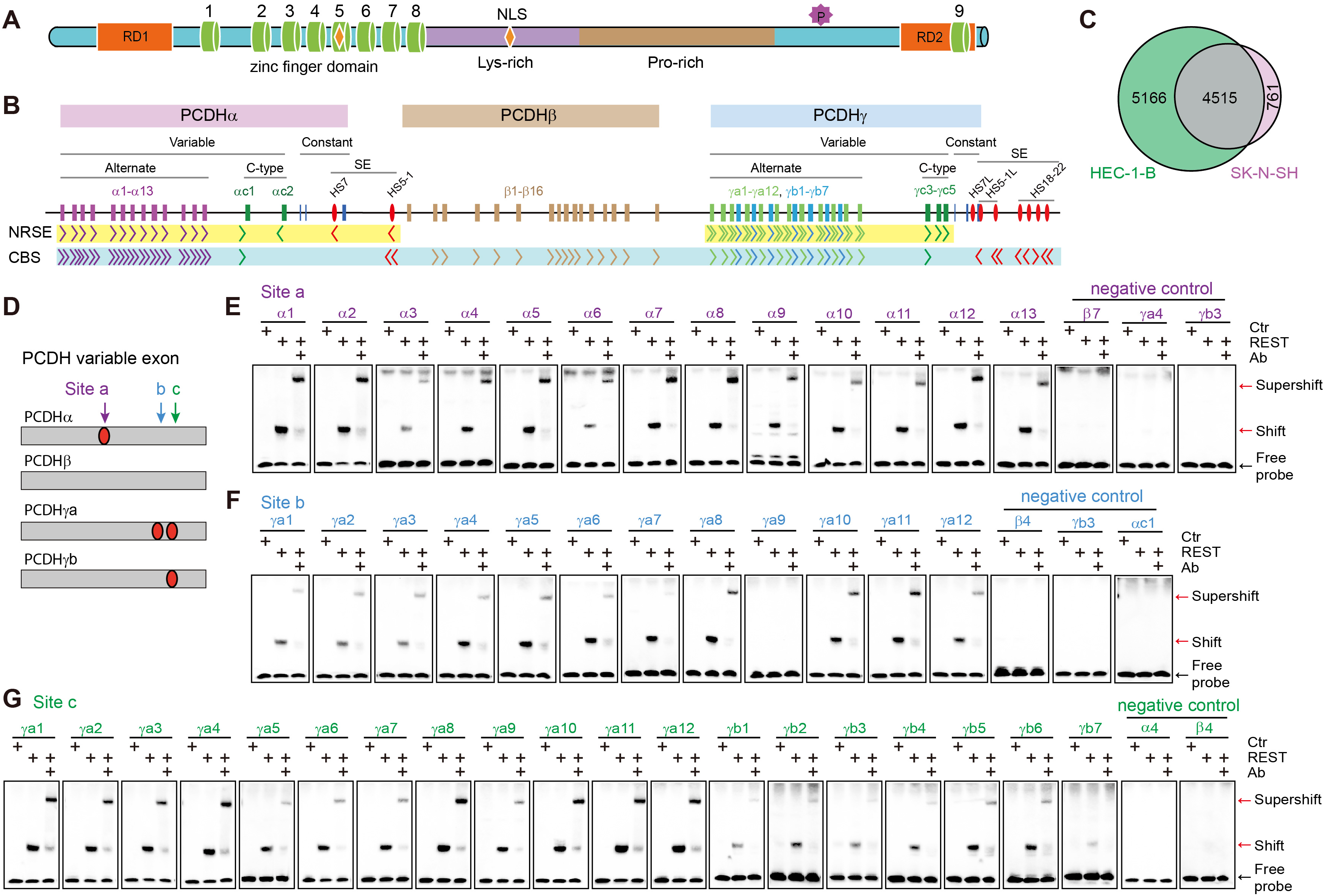
REST/NRSF binding sites in the human PCDH clusters. (**A**) Schematic of the REST/NRSF protein. REST/NRSF protein contains a DNA binding domain of eight ZFs (cylinders), two repression domains (RD1 and RD2), lysine-rich and proline-rich domains, two nuclear localization signals (NLS, shown in orange rhombus), a phosphodegron, and a C-terminal ZF domain. (**B**) Schematic of the three human *PCDH* gene clusters. The *PCDH α* and *γ* clusters share similar organization, with dozens of variable exons each of which is spliced to a single set of downstream constant exons. The variable exons are divided into alternate and C-type groups. The *PCDHβ* cluster contains only variable exons. The super-enhancer of the *PCDHα* cluster (red ellipses) is located between *PCDH α* and *β* clusters. The super-enhancer of the *PCDHβγ* clusters (red ellipses) is located downstream of *PCDHγ*. The locations and orientations of REST/NRSF sites (NRSE) and CTCF sites (CBS) are shown under the genes. (**C**) Venn diagram of REST/NRSF ChIP-nexus peaks of HEC-1-B and SK-N-SH cells. (**D**) Schematic of the NRSE locations within the four subgroups of the clustered *PCDH* variable exons. (**E**-**G**) EMSA experiments using REST/NRSF with a site ‘a’ NRSE probe of each member of the alternate *PCDHα* (**E**), a site ‘b’ NRSE probe of each *PCDHγa* (**F**), or a site ‘c’ NRSE probe of each member of the alternate *PCDHγ* (**G**), using mock as a control (Ctr). Supershifted bands were detected with a specific antibody against human MYC (Ab) tag fused to the C-terminal of REST/NRSF.

The clustered protocadherin (*PCDH*) genes encode a large number of cell-surface cadherin-like adhesion proteins which are thought to function as neuron identity codes in brain wiring and neuron discrimination (22–25). There are 53 highly-similar clustered *PCDH* genes organized into three closely-linked gene clusters (*PCDHα*, *PCDHβ*, and *PCDHγ*) in the human 5q31 chromosomal region (Figure 1B) (22). The genomic organizations of *PCDHα* and *PCDHγ* are similar in that each contains more than a dozen variable exons that can be spliced to a single set of three downstream constant exons. The variable exons can be grouped into alternate and C-type exons based on their locations and similarities (Figure 1B) (22). The *PCDHβ* cluster contains only variable exons but with no constant exon (22). Each variable exon has its own promoter (26, 27). Two super-enhancers, one composed of *HS7* (DNase I hypersensitive site 7) and *HS5-1*, the other composed of *HS7L* (HS7 like), *HS5-1L* (HS5-1like) and *HS18-22*, regulate the expression of the *PCDH α* and *βγ* clusters, respectively (Figure 1B) (25,28–33).

CTCF (CCCTC-binding factor) is a master regulator of clustered *PCDH* genes (25,31,33–37). There are tandem arrays of forward CTCF-binding sites (CBSs) associated with the *PCDH* promoters and of reverse CBSs associated with the super-enhancers (Figure 1B) (31, 33). Through CTCF/cohesin-mediated topological chromatin interactions between enhancers and promoters, clustered *PCDH* genes are stochastically and unbiasedly expressed (33, 37). By contrast, REST/NRSF has been shown to repress expression of the clustered *PCDH* genes (29, 38). However, the mechanism by which REST/NRSF represses these clustered genes remains unknown.

Here we identified NRSEs in every alternate exon of the human *PCDH α* and *γ* clusters, as well as in each C-type gene. By systematic EMSA (electrophoretic mobility shift assay) experiments, in conjunction with computational molecular dynamics, we found that REST/NRSF recognizes NRSEs via tandem ZF domains in a directional and flexible manner. Moreover, by genetic experiments we found that REST/NRSF inhibits long-distance chromatin contacts between the *PCDHα HS5-1* enhancer and its target promoters through preventing CTCF binding. Thus, REST/NRSF regulates the neural-specific expression of the clustered *PCDHα* genes through modifying chromatin structures.

## RESULTS

### A repertoire of NRSEs in clustered PCDH genes

To investigate mechanisms of neural-specific expression of the clustered *PCDH* genes in the brain, we performed REST/NRSF ChIP-nexus experiments in model cells of neuroblastoma SK-N-SH and endometrial carcinoma HEC-1-B (33,35,37) and found 5,276 and 9,681 REST/NRSF sites, respectively (Figure 1C). In the clustered *PCDH* loci, we found a repertoire of numerous REST/NRSF sites within variable regions of the *PCDH α* and *γ*, but not *β*, clusters (Figure 1B, Supplementary Figure S1A). In particular, there are two REST/NRSF sites in the variable exons of *PCDHγa* (Supplementary Figure S1B). Sequence analyses identified three conserved REST/NRSF sites, designated as ‘a’, ‘b’, and ‘c’, within the *PCDH* variable exons (Figure 1D, Supplementary Figures S2 and S3). Site ‘a’ in *PCDHα*, site ‘b’ in *PCDHγa*, and site ‘c’ in the *PCDHγ* alternate exons match the REST/NRSF consensus motif (Supplementary Figures S2 and S3) (16).

To investigate recognition mechanisms of these *PCDH* sites, we performed comprehensive EMSA experiments using recombinant REST/NRSF proteins. For site ‘a’, we designed a repertoire of probes for each alternate member of the *PCDHα* cluster and three probes in the same location for members of the *PCDHβγ* clusters as negative controls. EMSA experiments revealed that site ‘a’ in the *PCDHα* cluster, but not in the *PCDH β* or *γ* cluster, is recognized by REST/NRSF *in vitro*. Specifically, there is a shifted band with probes for each *PCDHα* alternate exon and a supershifted band when incubated with a specific antibody (Figure 1E). Similarly, for site ‘b’, only members of the *PCDHγa* subfamily (except *PCDHγa9*) are recognized by REST/NRSF (Figure 1F). Consistent with the nonbinding to the *PCDHγa9* probe, the first and fourth positions of the *PCDHγa9* site had been mutated from consensus ‘G’ and ‘C’ to non-consensus ‘A’ and ‘T’ nucleotides, respectively (Supplementary Figure S3A). For site ‘c’, members of both the *PCDHγa* and *PCDHγb* subfamilies are recognized by REST/NRSF *in vitro* (Figure 1G). These data suggest that there is one NRSE in each alternate exon of *PCDHα* (site ‘a’) and *PCDHγb* (site ‘c’), two NRSEs in each alternate exon of *PCDHγa* (sites ‘b’ and ‘c’, except *PCDHγa9*), and no NRSE in the exons of the *PCDHβ* cluster.

The five C-type *PCDH* exons are distinct from alternate exons both in their encoded protein sequences and regulatory mechanisms (22,35,37,39,40). We performed EMSA experiments for each member of the C-type exons and found that each contains a REST/NRSF site (Supplementary Figure S4A and B). We also systematically examined candidate recognition sites in the super-enhancers of the *PCDH α* and *βγ* clusters by EMSA experiments and found that *HS7* and *HS5-1* each contains a REST/NRSF site (Supplementary Figure S4C and D). However, in the super-enhancer of the *PCDHβγ* clusters, we could not find any authentic REST/NRSF site (Supplementary Figure S4C and E).

In summary, in the *PCDHα* cluster, both variable exons and the downstream super-enhancer contain NRSEs; but in the *PCDHβγ* clusters, only *PCDHγ* but neither *PCDHβ* nor their downstream super-enhancer contains NRSE, consistent with the neural-specific expression of *PCDHα*, but not *PCDH β* or *γ* genes (Supplementary Figure S4F) (41).

### Directional REST/NRSF binding to PCDH NRSEs

To investigate REST/NRSF DNA-recognition mechanisms, we generated a set of truncated REST/NRSFs by sequential deletion of ZF domains from either N- or C-terminus (Figure 2A-C). We found that, for the series of C-terminal deletion mutants, truncation up to ZF6 (ZF1-5) does not perturb REST/NRSF binding, but truncation up to ZF5 (ZF1-4) abolishes REST/NRSF binding to the noncanonical *HS5-1* site, suggesting that ZF5 plays an important role in the recognition of the *HS5-1* site (Figure 2D). The same result was also observed using the right-half only site of *PCDHα8*, suggesting that ZF5 is also essential for the recognition of the right-half site (Figure 2E). For the series of N-terminal deletion mutants, truncation up to ZF4 (ZF5-8) does not perturb the REST/NRSF binding to the noncanonical *HS5-1* site (Figure 2D), but remarkably results in a significant decrease of binding levels to the right-half only site of *PCDHα8* (Figure 2E), suggesting that ZF4 probably recognizes nucleotides within the right-half site. This is consistent with previous studies on a C-terminal truncated form of REST/NRSF (42). Finally, we tested the binding ability of ZF-truncated REST/NRSF to two canonical sites and found that ZF5 is also required for their binding (Supplementary Figure S5A).

**Figure 2.**
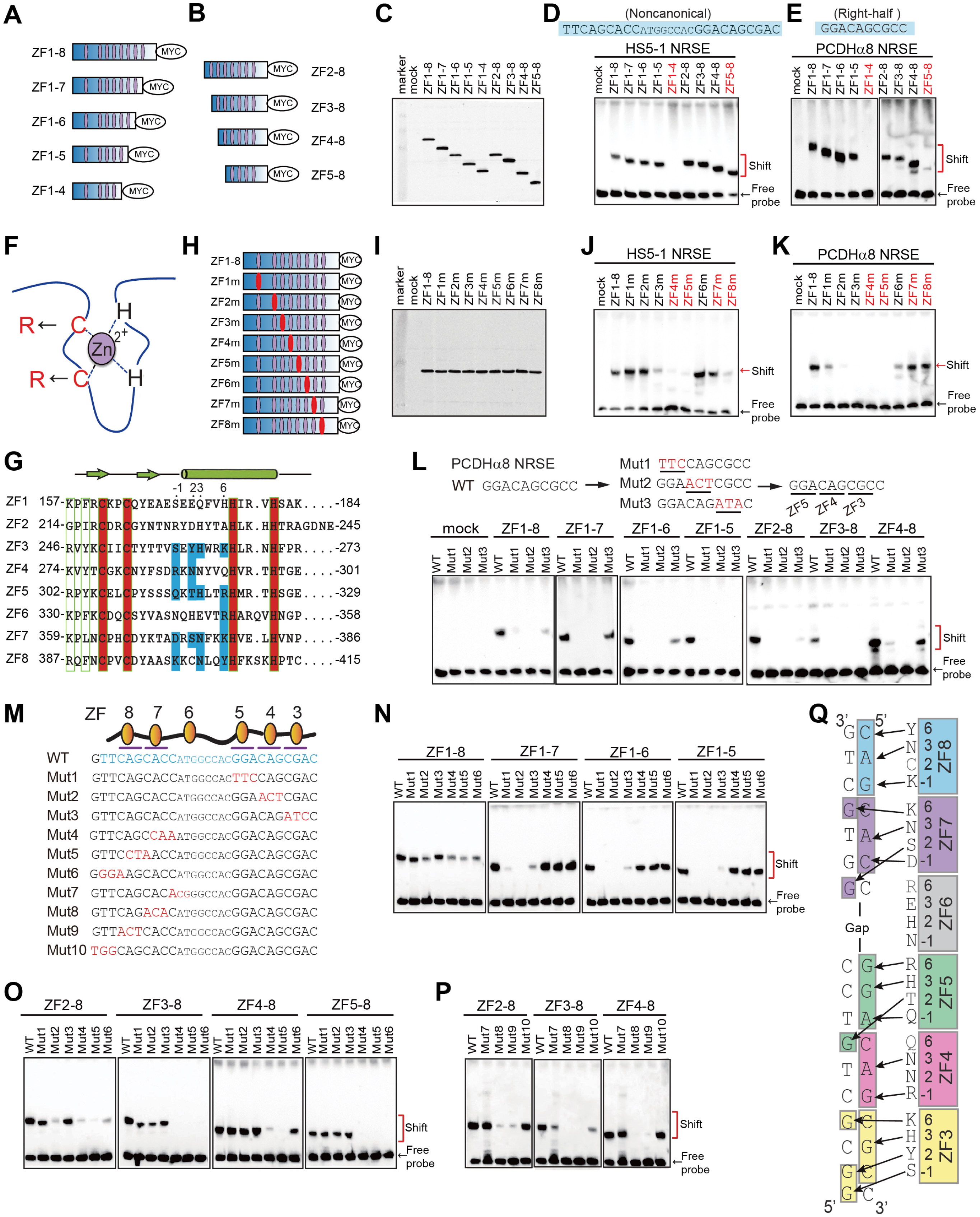
Directional base-specific binding of REST/NRSF to the clustered PCDH NRSEs. (**A**, **B**) Schematics of truncated REST/NRSF, fusing with a MYC tag, with sequential deletions of ZFs from either N or C terminus. Purple ellipses denote zinc-finger domains. (**C**) Western blot of REST/NRSF with sequential deletions of ZFs. (**D**, **E**) EMSA using the truncated proteins with biotinylated probes of noncanonical *HS5-1* NRSE (**D**) or of the right-half only *PCDHα8* NRSE (**E**). The sequences of the motifs are shown above the gel. (**F**) Schematic of point mutations in each ZF domain. Two key cysteine residues were mutated to two arginine residues. (**G**) Sequence alignment of the eight tandem Cys2-His2 zinc fingers of REST/NRSF, with the residues involved in chelating Zn^2+^ highlighted in the red background and residues involved in base contacts with NRSE motifs highlighted in the blue background. The −1, 2, 3, and 6 positions of the α-helix of each ZF domain are indicated above the sequences. (**H**) Diagram of the REST/NRSF mutants in each ZF domain (highlighted by red ellipses). **(I)** Western blot of ZF-mutated proteins. (**J**, **K**) EMSA using ZF mutated proteins with a biotinylated probe of the noncanonical *HS5-1* NRSE motif (**J**) or of the right-half only *PCDHα8* NRSE motif (**K**). (**L**) EMSA of the wild-type (WT) *PCDHα8* NRSE and its mutants using a set of ZF-deleted REST/NRSFs. (**M**) Sequences of the *HS5-1* NRSE WT (highlighted by blue) and mutant highlighted by red) motifs. (**N**-**P**) EMSA for ZF-deleted proteins using a biotinylated probe of the *HS5-1* NRSE Mut1-Mut6 (**N**, **O**) or Mut7-Mut10 (**P**), with the WT probe as a control. (**Q**) A model of base-specific contacts of ZF3-8 with DNA duplexes.

To further investigate the differential roles of ZF domains in NRSE recognition, we substituted the two Cys residues of each REST/NRSF tandem ZF domain with two Arg residues (Figure 2F-I). Substitution of Cys in ZF1 or ZF2 with Arg residues results in a significant decrease of the REST/NRSF binding to the right-half only site of *PCDHα8*, but does not perturb the binding to the noncanonical and canonical NRSEs (Figure 2J and K, Supplementary Figure S5B), suggesting that ZF1 and ZF2 are required for the binding to the right-half only site but are dispensable for REST/NRSF binding to the noncanonical and canonical NRSEs with both left- and right-half sites. In addition, substitution of Cys in ZF3, ZF4, or ZF5 with Arg residues affects the REST/NRSF binding to all NRSEs tested, including noncanonical, canonical, and right-half only sites (Figure 2J and K, Supplementary Figure S5B). This suggests that that ZF3-5 recognize the right-half sites. Finally, substitution of Cys in ZF7 or ZF8 with Arg residues does not affect REST/NRSF binding to the right-half only site of *PCDHα8*, but results in a significantly decrease of REST/NRSF binding levels to the noncanonical and canonical NRSEs (Figure 2J and K, Supplementary Figure S5B), suggesting that ZF7-8 recognize nucleotides within the left-half site. Together, these REST/NRSF truncation and mutation experiments suggest that ZF3-5 recognize the right-half while ZF7-8 recognize the left-half sites.

To further investigate the directionality of REST/NRSF DNA recognition, we sequentially mutated triple nucleotides (Mut1, Mut2 and Mut3) of the right-half only site of *PCDHα8* and found that these mutations abolish the binding of the C-terminal truncated REST/NRSF (Figure 2L). For the N-terminal truncated proteins, Mut3 abolishes the binding of ZF2-8 and ZF3-8, but not ZF4-8, suggesting that ZF3 binds to the three nucleotides corresponding to Mut3 (Figure 2L). Considering the essential role of ZF4 and ZF5 in binding to the right-half site (Figure 2D and E, J and K), we conclude that they recognize the triple nucleotides corresponding to Mut2 and Mut1, respectively.

We next generated a series of mutated NRSE probes of the *PCDH HS5-1* noncanonical site with sequential substitution of triple nucleotides (Figure 2M) and performed comprehensive EMSA experiments using REST/NRSF with sequential ZF deletions. Similar to the right-half only site, EMSA experiments demonstrated that mutations of the right-half (Mut1-3), but not left-half (Mut4-6), of the noncanonical site affect the REST/NRSF binding only if it contains ZF1-5 (Figure 2N). By contrast, mutations of the left-half (Mut4-6), but not right-half (Mut1-3), of the noncanonical site appear to affect the REST/NRSF binding only if it contains ZF5-8 (Figure 2O), again suggesting that ZF3-5 bind to the right-half site while ZF7-8 bind to the left-half site of NRSEs. Moreover, we generated an additional series of triple nucleotide mutations of the left-half site with a different phase (Mut7-10) (Figure 2M). Remarkably, we found that Mut8 and Mut9 almost abolish the REST/NRSF binding completely but Mut7 and Mut10 have much less effects (Figure 2P), suggesting that the six nucleotides corresponding to Mut8 and Mut9 play a major role. In conjunction with the fact that ZF6 mutation does not appear to affect the REST/NRSF binding (Figure 2J, Supplementary Figure S5B), we conclude that the six conserved nucleotides of the left-half site are recognized by ZF7-8. Finally, we mutated sequences downstream of the Mut3 site (Mut11 and Mut12) and found that they do not affect REST/NRSF binding, suggesting that ZF2 and ZF1 have no major role in binding the right-half only site of *PCDH*α8 (Supplementary Figure S5C). Taken together, we conclude that REST/NRSF base-specifically recognizes DNA duplexes in an antiparallel manner via tandem ZF3-8 (Figure 2Q).

### ChIP-nexus peaks with multiple NRSE motifs

We previously found that some CTCF peaks contains more than one CBS element (43). We also noted that some REST/NRSF peaks contain more than one NRSE. For example, one peak in the first exon of the neural protocadherin *CELSR3* gene contains four tandem half-site motifs in the configuration of left-11bp-right-9bp-left-11bp-right. These four tandem half-site motifs either function as four half-site only NRSEs or as two noncanonical NRSEs separated by 9 bp (Figure 3A). These sites have previously been shown to play an important role in the regulation of neural expression of the protocadherin *CELSR3* gene (44).

**Figure 3.**
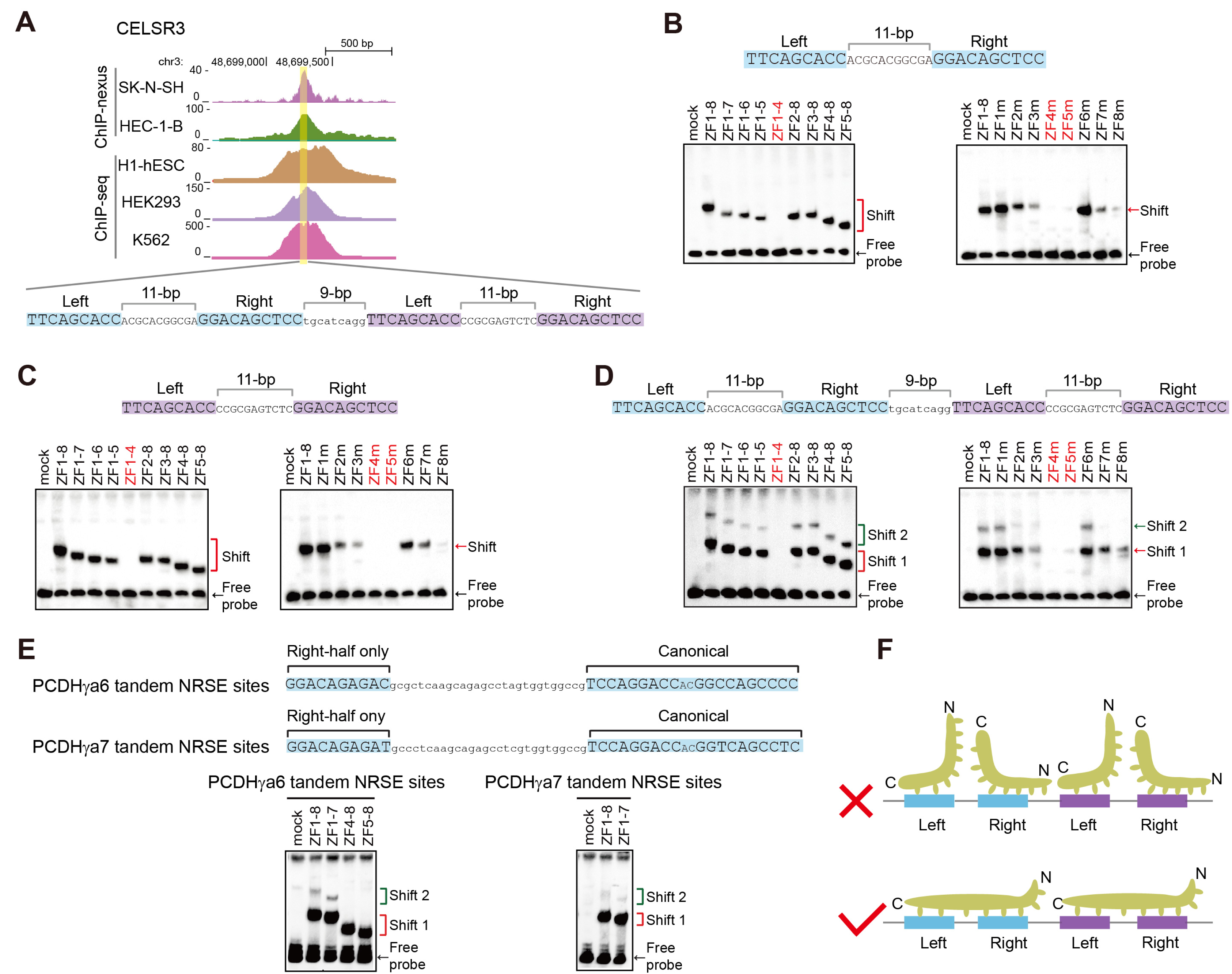
Recognition of tandem NRSE motifs by REST/NRSF. (**A**) Comparison of the REST/NRSF ChIP-nexus and ChIP-seq data of the human protocadherin *CELSR3* gene containing four tandem NRSE motifs. (**B**-**D**) EMSA for probes of the first two NRSE motifs (**B**), the last two NRSE motifs (**C**), or all of the four tandem NRSE motifs (**D**) with a repertoire of truncated or mutated REST/NRSFs, using WT as a control. (**E**) EMSA for probes of the *PCDHγa6* or *PCDHγa7* tandem NRSE motifs with truncated REST/NRSFs, using WT as a control. (**F**) Schematic of the stereo-hindrance and proper configuration of REST/NRSF recognition of tandem NRSE motifs.

To investigate how REST/NRSF binds to these composite tandem sites, we prepared two probes each matching one of the hypothetical noncanonical NRSEs as well as a probe containing all of the four tandem sites (Figure 3B-D). EMSA experiments showed only a single shifted band for either left or right noncanonical NRSE (Figure 3B and C); by contrast, we observed two shifted bands for the probe containing all four tandem sites (Figure 3D). Similar two shifted bands were also observed for the *PCDHγa* tandem site (Figure 3E, Supplementary Figure S3). Taken together, these observations suggest that some REST/NRSF ChIP-nexus peaks contain more than one NRSE in tandem and each NRSE is bound by one REST/NRSF protein molecule in only one orientation (Figure 3F).

### Distinct ZF6 conformations for canonical and noncanonical NRSEs

Analyzing REST/NRSF binding sites of ChIP-nexus data revealed that the gaps between left- and right-half sites are either 2 bp in canonical NRSEs or 7-9 bp in noncanonical NRSEs, but intriguingly could not be 3-6 bp (Figure 4A). To investigate how REST/NRSF tolerates the flexible length of gaps in NRSE motifs, we performed molecular dynamics (MD) experiments using ZF3-8 and the site ‘c’ canonical NRSE motif of *PCDHγa6* with a 2-bp gap. We found that these six ZFs wrap around the DNA duplex with the N-terminus of each ZF α-helix inserted into the major groove and this binding complex can keep stable during the MD simulations (Figure 4B), even though EMSA experiments demonstrated that ZF6 is not essential for binding to canonical NRSEs (Supplementary Figure S5A and B). We then performed MD experiments using ZF3-8 and the noncanonical NRSE motif of *HS5-1* with an 8-bp gap. We found that ZF3-5 and ZF7-8 wrap around the DNA duplex in the major groove and form base-specific contacts; however, in this case ZF6 flips out of the DNA major groove and remains parallel to the axis of DNA during the whole MD simulations to span the 8-bp gap (Figure 4C). Thus, REST/NRSF recognizes canonical and noncanonical NRSEs with remarkable conformation changes.

**Figure 4.**
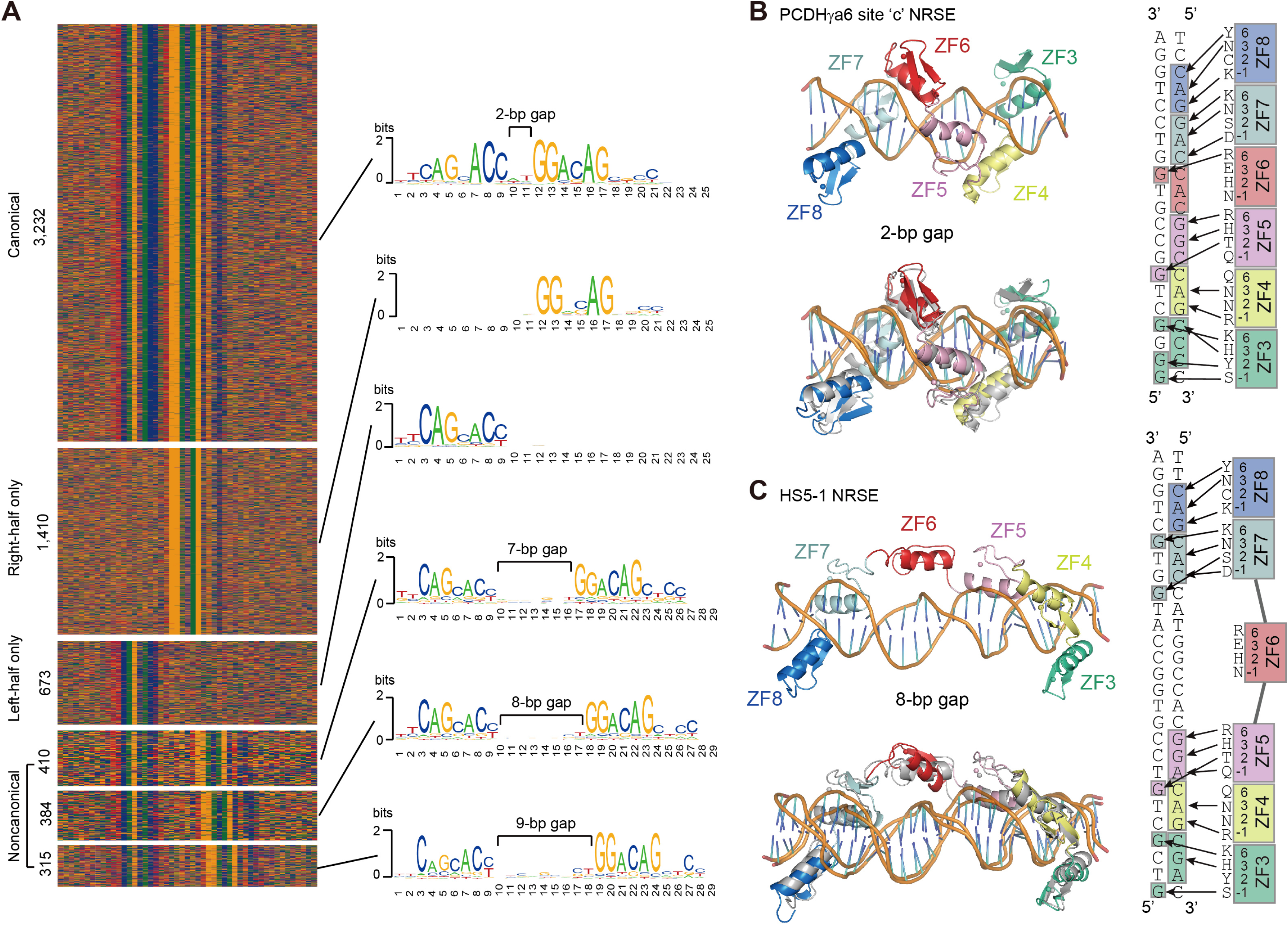
Flexible ZF6 determines two binding models of REST/NRSF with Canonical and Noncanonical NRSEs. (**A**) Six types of NRSEs and their motifs revealed by ChIP-nexus. (**B**, **C**) The molecular dynamics (MD) simulations for the base-specific contacts of ZF3-8 of REST/NRSF with site ‘c’ canonical NRSE of *PCDHγa6* with 2-bp gaps (**B**), or with *HS5-1* noncanonical NRSE with 8-bp gaps (**C**). In both (B) and (C), the upper panel is the initial structure of the MD simulations while the lower panel is the comparison of the initial and final structures. Base-specific contacts of ZF3-8 with the two NRSEs are shown on the right side of the panels.

We noted that in the 3,232 canonical NRSE motifs with 2-bp gaps, the 9^th^ position is a highly conserved base of ‘C’; however, this ‘C’ is barely conserved in the noncanonical NRSE motifs with variable large gaps of 7-9 bp (Figure 4A). Nevertheless, our EMSA experiments demonstrated that ZF7-8 and ZF3-5 contact with the left- and right-half sites, respectively, of both canonical and noncanonical NRSEs in a base-specific manner (Figures 1-3, Supplementary Figure S5). To investigate why the 9^th^ position ‘C’ is more conserved in canonical than noncanonical NRSEs, we simulated base-recognition of ZF6 by MD in the two different conformations. We found that, in the canonical NRSEs, the side chain of Arg349 in ZF6 can form stable hydrogen bonds with the base ‘G’ in the complementary strand (Figure 4B, Supplementary Figure S5D). In the noncanonical NRSEs, however, ZF6 is not a reader of any NRSE base but is positioned parallel to the DNA axis to span the much larger gap (Figure 4C). This explains that the 9^th^ position ‘C’ of canonical NRSEs is much more conserved than that of the noncanonical NRSEs (Figure 4A).

### REST/NRSF inhibits long-distance enhancer-promoter contacts

Most of NRSEs are located distal from promoters (16, 17). To investigate how these NRSEs regulate gene expression, we focused on the *HS5-1* noncanonical site and perturbed either the REST/NRSF protein or the NRSE *cis*-element. We first knocked down REST/NRSF by designing three different shRNAs and found that REST/NRSF knockdown results in a significant increase of the expression levels of *PCDH α6* and *α12* genes in HEC-1-B cells (Figure 5A and B). In addition, we found that deposition of active chromatin marks, H3K4me3 and H3K27ac, is also significantly increased in both the *HS5-1* enhancer and *PCDHα* promoters (Figure 5C-E), suggesting that REST/NRSF regulates *PCDHα* expression through epigenetic modifications. Because the expression of *PCDHα* genes depends on their long-distance chromatin interactions with the distal enhancer (35), to see whether the 3D chromatin architecture of the locus is altered, we performed quantitative high-resolution chromosome conformation capture followed by next-generation sequencing (QHR-4C) experiments (33) with either *HS5-1* or *PCDHα12* as a viewpoint. Remarkably, we found that there is a significant increase of long-distance chromatin interactions between the *HS5-1* enhancer and its target *PCDHα* genes upon REST/NRSF knockdown (Figure 5F and G), which explains the increased expression levels of the *PCDHα* genes in HEC-1-B cells (Figure 5B). Similar effects were also observed in HEK293T cells upon REST/NRSF knockdown (Supplementary Figure S6A-C).

**Figure 5.**
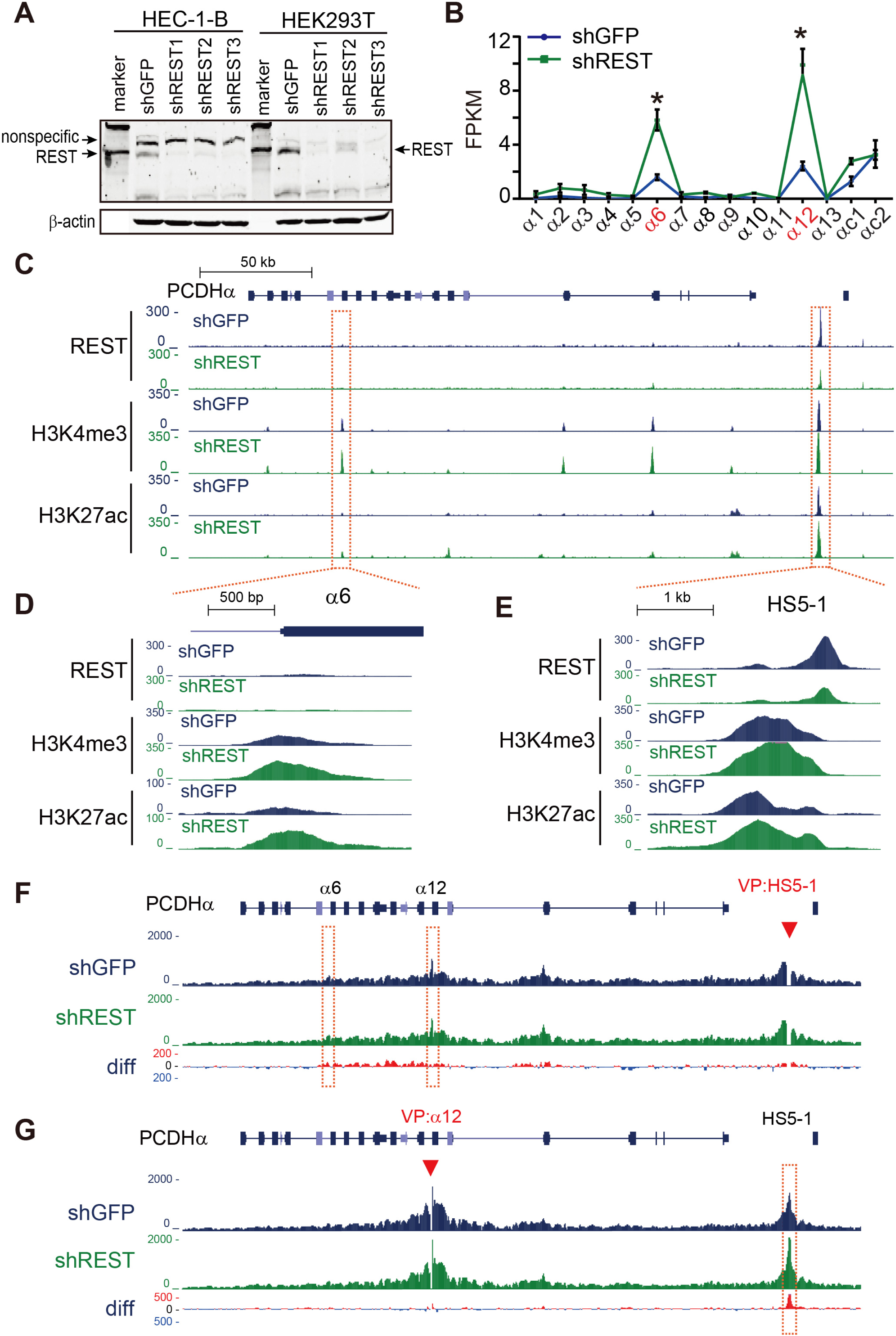
REST/NRSF knockdown enhances *HS5-1* activity and long-distance enhancer-promoter contacts. (**A**) Western blot showing the REST/NRSF knockdown efficiency of the three shRNAs in HEC-1-B and HEK293T cells. (**B**) RNA-seq showing the increased *PCDHα* expression upon REST/NRSF knockdown in HEC-1-B cells. Data are presented as mean ± SEM. * *P < 0.05*. (**C**-**E**) ChIP-seq with antibodies against REST/NRSF, H3K4me3, and H3K27ac in control and REST/NRSF knockdown cells. Note the decrease of REST/NRSF and the increase of H3K4me3 and H3K27ac upon REST/NRSF knockdown. (**F**, **G**) QHR-4C interaction profiles of the *PCDHα* locus using *HS5-1* (**F**), or *PCDHα12* (**G**) as a viewpoint (VP, arrowheads) for the control and REST/NRSF knockdown cells. Differences (shREST vs shGFP) are shown under the 4C profiles.

We then deleted the NRSE within the *HS5-1* enhancer in HEC-1-B and HEK293T cells by screening single-cell CRISPR clones by DNA-fragment editing (45, 46). We obtained two single-cell homozygous clones with NRSE deletion in each cell line (Supplementary Figure S6D-G). RNA-seq revealed that deletion of the *HS5-1* NRSE results in a significant increase of expression levels of the *PCDH α6* and *α12* genes (Figure 6A, Supplementary Figure S6H). In addition, the deposition of H3K4me3 and H3K27ac histone marks is also significantly increased in both the *HS5-1* enhancer and *PCDHα* target promoters in these single-cell CRISPR clones (Figure 6B-D, Supplementary Figure S6I-K). Similar to the REST/NRSF knockdown, deletion of the *HS5-1* NRSE also results in a significant increase of long-distance chromatin interactions between the *HS5-1* enhancer and *PCDHα* target promoters (Figure 6E and F). Together, these data suggest that *HS5-1*-bound REST/NRSF suppresses the expression of *PCDHα* genes through modulation of histone tails and long-distance chromatin interactions.

**Figure 6.**
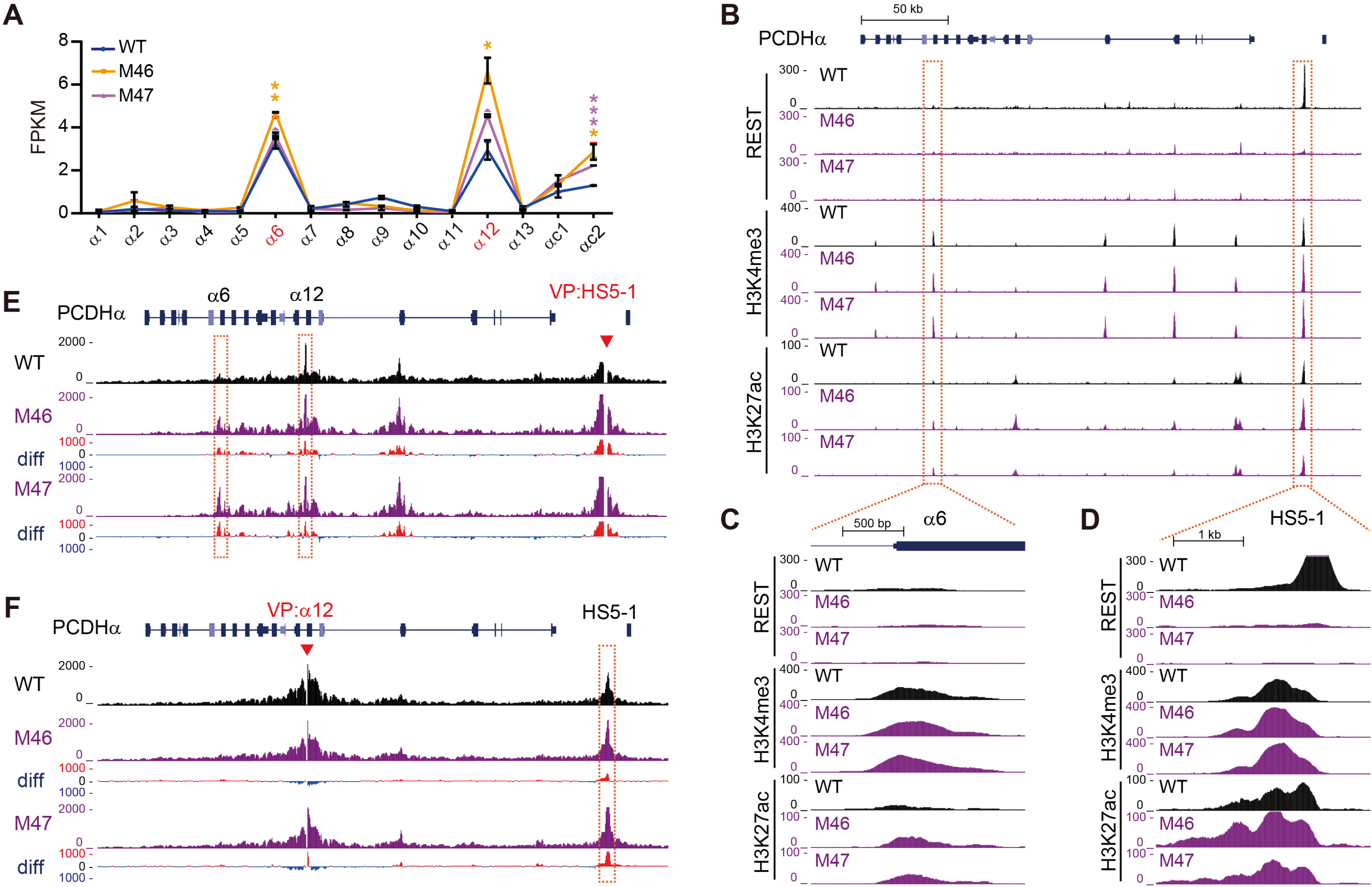
*HS5-1* NRSE deletion results in increased *HS5-1* enhancer activity and long-distance enhancer-promoter contacts. (**A**) RNA-seq results showing the increased *PCDH α6* and*α12* expression upon *HS5-1* NRSE deletion in HEC-1-B cells. M46 and M47 are two CRISPR single-cell clones with *HS5-1* NRSE deletion. Data are presented as mean ± SEM. * *P < 0.05*, *\*\* P < 0.01, *** P < 0.001*. (**B**-**D**) ChIP-seq with a specific antibody against REST/NRSF, H3K4me3, or H3K27ac in WT and *HS5-1* NRSE deletion cells. (**E**, **F**) QHR-4C interaction profiles of the *PCDHα* locus using *HS5-1* (**E**), or *PCDHα*12 (**F**) as a viewpoint (VP, arrowheads) for the WT and *HS5-1* NRSE deletion single-cell clones. Differences (deletion vs WT) are shown under the 4C profiles.

### NRSE deletion reshapes Pcdhα 3D chromatin structure in vivo

To see whether *HS5-1*-bound REST/NRSF modulates histone tails and chromatin architectures *in vivo*, we deleted the *HS5-1 NRSE* in mice by CRISPR pronuclear injection with dual sgRNAs (Figure 7A, Supplementary Figure S7A and B). ChIP-seq showed that the binding of REST/NRSF to *HS5-1* is abolished upon NRSE deletion (Supplementary Figure S7C). We first confirmed that there are significant higher levels of REST/NRSF expression in kidney than in cortical tissues by Western blot and RNA-seq (Figure 7B and C). Interestingly, there is a significant increase of expression levels of members of the *Pcdhα* cluster in kidney but not in cortical tissues upon deletion of the *HS5-1* NRSE (Figure 7D and E). In addition, the deposition of active histone mark of H3K4me3 is significantly increased in *HS5-1* and in members of the *Pcdh*α cluster but not in *Pcdhβ1* in kidney tissues (Figure 7F).

**Figure 7.**
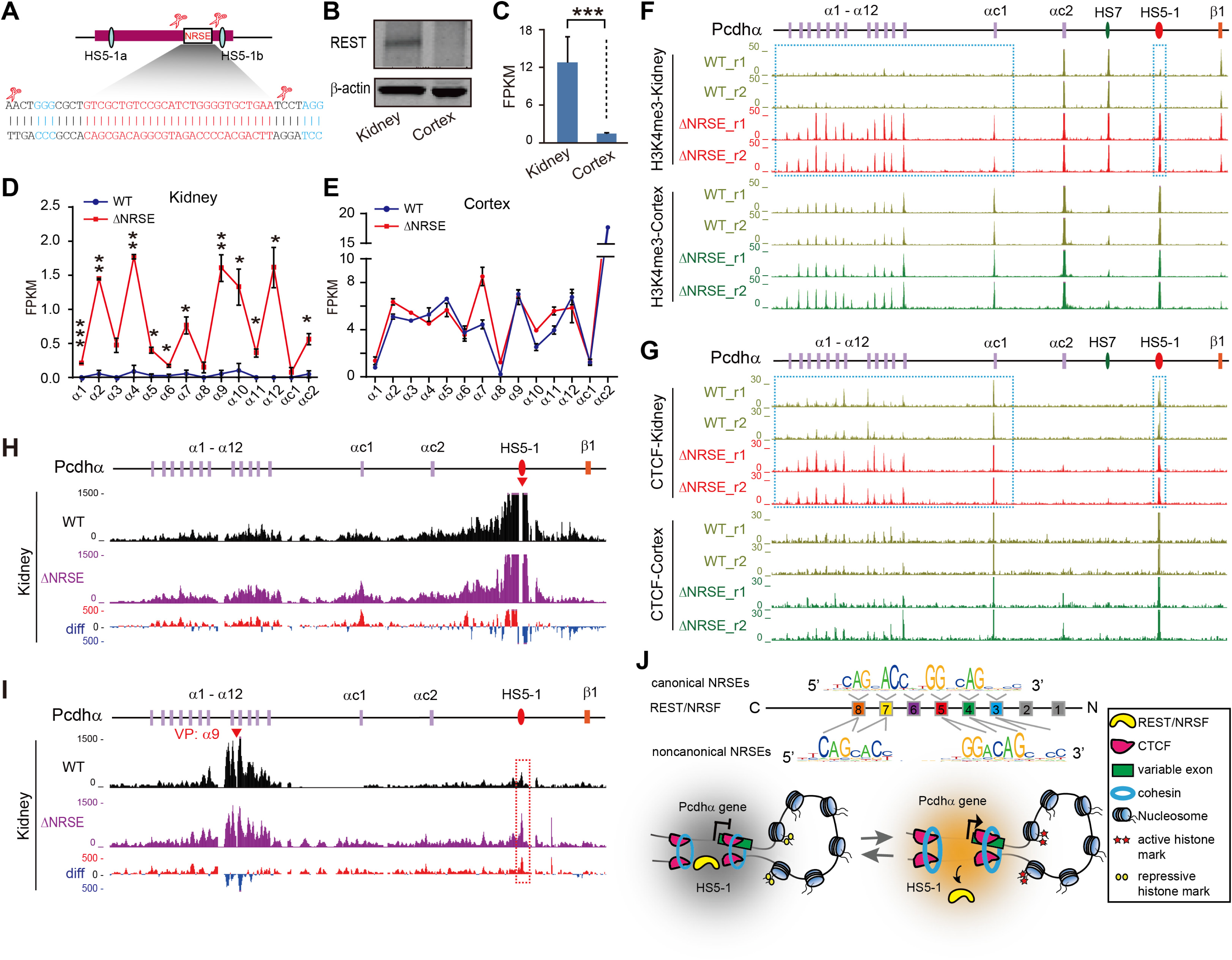
NRSE deletion reshapes the *Pcdhα* 3D chromatin structure in mice *in vivo* via CTCF enrichments. (**A**) Schematic of the *HS5-1* enhancer showing the precise deletion of NRSE by CRISPR DNA-fragment editing with dual sgRNAs. The PAM sites are highlighted. The two CTCF binding sites (HS5-1a and HS5-1b) are marked with cyan ovals. (**B**, **C**) The expression levels of REST/NRSF in the kidney and cortical tissues of mice tested by Western blot (**B**) or RNA-seq (**C**). (**D**, **E**) RNA-seq showing the significant increase of the *Pcdha* expression levels in mouse kidney (**D**), but not in cortical (**E**), tissues upon NRSE deletion. Data are presented as mean ± SEM. *\* P < 0.05*, *\*\* P < 0.01, *** P < 0.001*. (**F**, **G**) ChIP-seq of H3K4me3 (**F**) and CTCF (**G**) in the *Pcdhα* cluster in the kidney and cortical tissues of WT and *HS5-1* NRSE-deleted (ΔNRSE) mice. Note the significant increases of H3K4me3 and CTCF in kidney upon NRSE deletion. (**H**, **I**) QHR-4C interaction profiles of the *Pcdhα* locus using *HS5-1* (**H**) or *PCDHα9* (**I**) as a viewpoint (VP, arrowheads) for the kidneys of WT and ΔNRSE mice. Differences (deletion vs wild-type) are shown under the 4C profiles. (**J**) A model depicting the DNA-recognition and mechanism of REST/NRSF repressing *PCDHa* gene expression through CTCF/cohesin-mediated higher-order chromatin structures.

To investigate why the distal sites were affected, we performed CTCF ChIP-seq experiments and found that there is a significant increase of CTCF enrichments in *HS5-1* and in members of the *Pcdh*α cluster in kidney tissues (Figure 7G, Supplementary Figure S7D and E). Finally, QHR-4C experiments demonstrated that there is a significant increase of long-distance chromatin interactions between the distal *HS5-1* enhancer and *Pcdhα* target genes upon NRSE deletion (Figure 7H and I). These data suggest that REST/NRSF represses expression of *Pcdhα* target genes through modulating higher-order chromatin structures in mice *in vivo* (Figure 7J).

## DISCUSSION

The clustered *PCDHs* participate in a wide variety of neurodevelopmental processes such as dendritic self-avoidance, axonal even spacing and tiling, spine morphogenesis and synaptogenesis, neuronal migration and survival (23, 47). Through stochastic and combinatorial expression, the clustered *PCDH* genes encode countless assemblies of cell-surface molecules to endow each neuron with a unique identify code (33,37,39,48,49). Thus, the spatiotemporal expression patterns of the clustered *PCDH* genes must be precisely regulated during brain development. REST/NRSF is a key repressor of neural genes in neuronal progenitors and its derepression is central for neurogenesis and neuronal differentiation (2,3,8). Here we found that REST/NRSF recognizes diverse *PCDH* NRSEs in an antiparallel manner via base-specific contacts with tandem ZF domains. In addition, MD simulations revealed that REST/NRSF endures different gap sizes of canonical and noncanonical NRSEs by adopting distinct conformations for ZF6. Finally, through genetic approaches, we demonstrated that the mechanism by which enhancer-bound REST/NRSF represses the expression of distant *PCDHα* genes is through modulating higher-order chromatin structures.

The stochastic expression of clustered *PCDHα* genes is achieved through promoter choice determined by CTCF/cohesin-mediated chromatin interactions between the *HS5-1* enhancer and its target promoters (29,31,35,37). In the *PCDHβγ* clusters, topological chromatin interactions between tandem CTCF sites determine the balanced expression of members of the *PCDHβγ* genes (33). In contrast, how *PCDH* genes are silenced in neural progenitors and nonneural tissues remains largely unknown. We showed here that binding of REST/NRSF to the *HS5-1* enhancer inhibits the expression of the *PCDHα* genes through CTCF/cohesin-mediated chromatin interactions. We identified NRSEs in each member of the *PCDH* α and γ gene clusters. These sites may be related to the repression of nonchosen members of clustered *PCDH* α or γ genes in single cells in the brain because REST/NRSF has been shown to directly repress neural gene expression (6,50,51).

REST/NRSF binds to diverse NRSEs *in vivo* with distinct and hierarchical affinities according to their sequence variations (16,52,53). It is known that ZF domains are important for REST/NRSF binding to DNA duplexes (16, 42); however, it is puzzling why there exist two major classes of canonical and noncanonical NRSEs with distinct gaps in the human genome (16). By a combination of EMSA and molecular dynamics experiments, we found that, in the case of canonical NRSEs, each ZF domain of tandem ZF3-8 is inserted into the major groove of DNA duplexes and recognizes standard 3 bp (Figure 4B). By contrast, in the case of noncanonical NRSEs, ZF7-8 and ZF3-5 are inserted into the major grooves of the left- and right-half sites. However, in this case, ZF6 is flipped out of the major groove and positioned nearly parallel to the axis of the DNA duplexes, serving as a spacer element to tolerate variable distances between the two half sites. Thus, ZF6 functions as bridge-like spacer for the flexible gaps connecting the left- and right-half sites of the noncanonical NRSE motifs (Figure 4C). This explains the long-standing mystery that REST/NRSF recognizes both canonical and noncanonical classes of NRSE motifs across the entire human genome (16).

MD demonstrated that, for the canonical NRSEs, the Arg349 of the ZF6 forms base-specific contacts with the ‘G’ base of the complementary strand at the 9^th^ position (Figure 4B, Supplementary Figure S5D), explaining the observed conservation of the ‘C’ nucleotide at the 9^th^ position of canonical NRSE motifs (Figure 4A). Moreover, for the noncanonical NRSEs, once ZF6 flipped out of the major groove, it cannot form base-specific contacts with any nucleobase. Instead, the flexible linkers between ZF5 and ZF6, and between ZF6 and ZF7 allow REST/NRSF to tolerate variable gap distances between the left- and right-half sites of noncanonical NRSE motifs (Figure 4C). Finally, because of the biophysical hindrance of the flipped ZF6, the gap distance between the left- and right-half sites cannot be 3-6 bp, consistent with the fact that the flexible gaps of the noncanonical NRSEs cannot be less than 7 bp in the human genome (Figure 4). Together, our data shed significant insights into mechanisms by which REST/NRSF recognizes diverse genomic DNA sites and regulates gene expression.

## MATERIALS AND METHODS

### Cell culture and transfection

HEK293T cells were cultured in Dulbecco’s modified Eagle’s medium (Hyclone) supplemented with 10% (v/v) FBS (Gibco), and 1% penicillin-streptomycin (Gibco). HEC-1-B and SK-N-SH cells were cultured in MEM (Hyclone), supplemented with 10% (v/v) FBS, 2 mM GlutaMAX (Gibco), 1 mM sodium pyruvate (Sigma), and 1% penicillin-streptomycin. Cells were maintained at 37 °C in a 5% (v/v) CO_2_ incubator. Cells were transfected by Lipofectamine 2000 (Invitrogen) when the cell confluency reached 70%∼90%.

### Animals

C57BL/6 and ICR mouse strains were housed at 23°C on a 12/12 h light-dark cycle (7:00 am–19:00 pm) in the SPF facilities. The zygotes were obtained from the oviducts of superovulated female C57BL/6 mice mated with the C57BL/6 stud male mice. All experiments were carried out in accordance with Institutional Animal Care and Use Committee of Shanghai Jiao Tong University (protocol#: 1602029).

### Plasmid construction

To generate plasmids for *in-vitro* expression of REST/NRSF, the coding sequence of REST/NRSF ZF1-8 were amplified by PCR using cDNA as templates and cloned into the pTNT vector (Promega) between the EcoRI and XbaI sites, with a MYC tag in the 3’ terminal. A series of ZF-deleted and ZF-mutated REST/NRSF-expressing plasmids were constructed by PCR using ZF1-8 as templates and ligated into the pTNT vector between the EcoRI and XbaI sites, also with a MYC tag in the 3’ terminal.

To generate plasmids used as probe templates for the EMSA experiments, the sequences containing putative NRSE motifs were amplified by PCR from the human genomic DNA and then subcloned into the pGEM-T Easy vector (Promega). To generate plasmid templates for the mutated probes, the mutated sequences were constructed by PCR from the wild-type plasmids and then ligated into the pGEM-T Easy vector.

The knockdown plasmids for REST/NRSF and for the control GFP were constructed by ligating annealed primer pairs into the pLKO.1 vector (Addgene) between the EcoRI and AgeI sites. Plasmids for sgRNA expression were constructed by inserting annealed primer pairs into a BsaI-linearized pGL3 vector under the control of the U6 promoter. All constructs were confirmed by sequencing and all primers are shown in Supplementary Table S1.

### Lentivirus packaging and infection

The pLKO.1-plasmids for REST/NRSF knockdown were co-transfected into HEK293T cells with the psPAX2 and pMD2.G helper plasmids (Addgene) using Lipofectamine 2000 (Life Technologies) to produce lentiviral particles. HEC-1-B cells and HEK293T cells at the confluency of 70%∼90% were infected with the virus in the presence of 8 μg/ml polybrene (Sigma). Puromycin (Sigma) was added at a final concentration of 2 μg/ml to select the infected cells. Fresh puromycin-containing medium was changed every other day. Cells were collected for assays at day 5 post infection. After harvested by lysing cells with RIPA lysis buffer (50mM Tris-HCl, pH 7.4, 150mM NaCl, 1% Triton X-100, 1% sodium deoxycholate, 0.1% SDS, 1 mM PMSF), the expression levels of REST/NRSF were measured by Western blot.

### Western blot

Proteins were denatured and separated by SDS–PAGE and transferred to nitrocellulose membranes. These membranes were then incubated with mouse anti-MYC (Millipore), rabbit anti-REST (Millipore), or rabbit anti-β-actin antibody (Abcam). Finally, the membranes were incubated with anti-mouse or anti-rabbit secondary antibodies and scanned using the Odyssey System (LI-COR Biosciences).

### Electrophoretic mobility shift assays (EMSA)

EMSA experiments were performed as described (31, 35) with some modifications. Proteins used for EMSA experiments were synthesized *in*-*vitro* from pTNT plasmids using TNT T7 Quick Coupled Transcription/Translation System (Promega) according to the manufacturer’s protocol. Briefly, ZF-deleted or ZF-mutated pTNT plasmids were mixed gently with TNT T7 Quick Master Mix, Methionine, and T7 TNT PCR Enhance by pipetting, and then incubated at 30 °C for 60–90 minutes.

Probes were generated by PCR with high-fidelity polymerase using 5’ biotin-labeled primers from template-containing plasmids and were gel-purified. The primers used are listed in Supplementary Table S1. Probe concentration was measured with NanoDrop (Thermo). Each binding reaction contained equimolar of the biotin-labeled probes. Protein concentration was determined by Western blot and each binding reaction contained the same amounts of proteins.

EMSA was performed with the LightShift Chemiluminescent EMSA Kit (Thermo) following the manufacturer’s manuals. Briefly, the *in-vitro*-synthesized proteins were precleared with the binding buffer containing 10 mM Tris-HCl, 250 mM KCl, 2.5 mM MgCl_2_, 0.1 mM ZnSO_4_, 1 mM DTT, 0.1% NP-40, 50 ng/μl poly (dI-dC), and 2.5% (v/v) glycerol on ice for 20 min. 50 fmol of biotin-labeled probes were then added and the reactions were incubated at room temperature for 20 min. 1μg of anti-MYC antibody was added into the binding reaction and incubated at room temperature for another 20 min for the supershift experiments. The binding reactions were electrophoresed on 5% nondenaturing polyacrylamide gels in ice-cold 0.5× TBE buffer (pH8.0) and transferred to a nylon membrane. After crosslinking under UV-light for 10 min, the membranes were blocked in Blocking Buffer by incubating for 15 min with gentle shaking and then incubated in Stabilized Streptavidin-Horseradish Peroxidase Conjugate solution for 15 min with gentle shaking. After rinsing the membrane with 1× washing buffer briefly, washed the membrane four times for 5 min each in 1× washing buffer with gentle shaking. Then, the membrane was incubated in substrate equilibration buffer for 5 min with gentle shaking, followed by incubation in the substrate working solution for 5 min without shaking and exposure using ChemiDoc XRS+ System (Bio-Rad).

### ChIP-nexus

ChIP-nexus experiments were performed as described (54) with some modifications. Briefly, ∼2 x 10^7^ cells were cross-linked with 1% formaldehyde for 10 min at room temperature, followed by quenching the crosslinking with glycine at a final concentration of 0.125 M and then spun down. Subsequently, cell pellets were lysed twice with ice-cold ChIP buffer (10 mM Tris-HCl, pH 7.5, 0.15 M NaCl, 1% Triton X-100, 1 mM EDTA, 0.1% SDS, 0.1% sodium deoxycholate, 1x protease inhibitors) by incubating at 4 °C for 10 min with slow rotation. Nuclei were spun down and then resuspended in 700 μl of the ChIP buffer. After incubating on ice for 10 min, the samples were sonicated using a Bioruptor Sonicator on high power (30 sec on/30 sec off) for 30 min to fragmentize DNA to sizes ranging from 100 to 10,000 bp. The sheared chromatin solutions were immunoprecipitated with a specific antibody against REST/NRSF (Millipore, 4 μg for each reaction) by slow rotation at 4 °C overnight. Antibody-precipitated complexes were incubated with 50 μl of Protein A/G Magnetic beads (Thermo) at 4 °C for another 3h the next day. The chromatin-enriched magnetic beads were then washed with the washing buffer A (10 mM TE, 0.1% Triton X-100), washing buffer B (150 mM NaCl, 20 mM Tris-HCl, pH 8.0, 5 mM EDTA, 5.2% sucrose, 1.0% Triton X-100, 0.2% SDS), washing buffer C (250 mM NaCl, 5 mM Tris-HCl, pH 8.0, 25 mM HEPES, 0.5% Triton X-100, 0.05% sodium deoxycholate, 0.5 mM EDTA), washing buffer D (250 mM LiCl, 0.5% NP-40, 10 mM Tris-HCl, pH 8.0, 0.5% sodium deoxycholate, 10 mM EDTA), and finally the Tris buffer (10 mM Tris-HCl, pH 7.5). Washing volumes were 1 ml per sample. After the washing buffer was added, the tubes were briefly inversed by hands to resuspend the beads for each washing. The beads were resuspended every ∼15 min by gently tapping the tubes for all the following incubations.

The DNA-complex-coated beads were incubated with the end-repair enzyme mixture (NEB) at 20 °C for 30 min to repair the DNA ends and then with Klenow exo- (NEB) in the NEB buffer 2 containing 0.2 mM dATP at 37 °C for 30 min for dA tailing. The samples were then ligated with the annealed Nexus adaptors (Nex_adaptor_UBamHI: 5’ phosphate GATCG GAAGA GCACA CGTCT GGATC CACGA CGCTC TTCC, Nex_adaptor_Barcode_BamHI: 5’ phosphate TCAGA GTCGA GATCG GAAGA GCGTC GTGGA TCCAG ACGTG TGCTC TTCCG ATCT) with 2x Blunt/TA ligase master mix (NEB) at 25 °C for 1h. The adaptors contain a pair of sequences for library amplification, a BamHI site for later linearization, a nine-nucleotide barcode containing five random bases and four fixed bases. Subsequently, the samples were treated with Klenow exo- at 37 °C for 30 min to fill the ends of adaptors and then trimmed with T4 DNA polymerase (NEB) at 12 °C for 5 min. The blunt-ended DNA was then treated with Lambda Exonuclease (NEB) at 37 °C for 60 min with constant rotation to digest one strand of the double-stranded DNA in a 5’ to 3′ direction until encountering a cross-linked protein. The samples were then digested by RecJ_f_ exonuclease (NEB) for 60 min at 37 °C to degrade the single-stranded DNA from the 5’-end and washed three times with the RIPA buffer (50 mM HEPES, pH 7.5, 1 mM EDTA, 0.7% sodium deoxycholate, 1% NP-40, 0.5 M LiCl).

The DNA-protein complex was eluted with 200 μl of elution buffer (50 mM Tris-HCl, pH 8.0, 10 mM EDTA, 1% SDS) at 65 °C, shaken at 1000 rpm for 30 min, and then reverse-crosslinked at 65 °C overnight. The next day, after incubating the samples with 2 μl of RNase A (Thermo) at 37 °C for 2h and then with 8 μl of proteinase K (NEB) at 55 °C for 4h, ssDNA was extracted with phenol: chloroform: isopentanol (25:24:1, v/v), and precipitated with 2.5x (v/v) of ethanol, 1/10 (v/v) of 3M sodium acetate and 1.5 μl of glycogen (20 mg/ml, Thermo). The DNA pellet was resuspended in 10 μl of nuclease-free water.

After denatured at 95 °C for 5 min, the ssDNA was circularized with CircLigase (Epicentre) at 60 °C for 1h. The samples were annealed with the BamHI_cut_oligo (GAAGA GCGTC GTGGA TCCAG ACGTG) in the digesting buffer (Thermo) in a thermocycler using the annealing program (95 °C for 1 min, slowly cooled down at 1% ramp to 25 °C for 1 min, 25 °C for 30 min, hold at 4 °C) and then digested with BamHI (Thermo) at 37 °C for 30 min. The linearized DNA were precipitated using ethanol and sodium acetate, then resuspended with 20 μl of water. The DNA was amplified by Q5 polymerase (NEB) with the Illumina primers with the PCR program (98 °C for 30 sec, 16 cycles of 98 °C for 10 sec and 65 °C for 75 sec, 65 °C for 5 min, hold at 4 °C). The PCR products with sizes from 150 bp to 300 bp were extracted from 2% agarose gel using MinElute Gel Extraction Kit (Qiagen).

ChIP-nexus library DNA was sequenced on an Illumina HiSeq X Ten platform, and reads were filtered for the presence of the fixed barcode CTGA starting from the 6^th^ position of reads. The random and fixed barcode sequences were then removed. Adaptor sequences at 3’ end of reads were then trimmed using the Cutadapt tool. The trimmed reads were aligned to hg19 using Bowtie2, and then analyzed by MACS2, MACE, and MEME suites. The heatmaps were generated by an R package named pheatmap.

### RNA-seq

RNA-seq experiments were performed as previously described (31, 33). Briefly, total RNA was extracted from cultured cells or mouse tissues using TRIzol Reagent (Ambion). RNA-seq experiments were performed using the NEBNext Ultra RNA Library Prep Kit for Illumina (NEB) according to the manufacturer’s protocol. Briefly, mRNA was enriched from 1 μg of total RNA using NEBNext oligo (dT) magnetic beads, and fragmented by heating at 95 °C for 10 min. After reverse transcription of the first stranded cDNA and synthesis of the second stranded cDNA, cDNA was purified using 1.8x AMPure XP beads (Beckman). The purified cDNA was end-repaired and then ligated with NEBNext Adaptors, followed by treatment with the USER enzyme (NEB). The ligated product was purified using the AMPure XP beads (Beckman). The purified cDNA product was then amplified with the Illumina primers by the Q5 enzyme (NEB) with the PCR program (98 °C for 30 sec, 14 cycles of 98 °C for 10 sec and 65 °C for 75 sec, 65 °C for 5 min, hold at 4 °C). RNA-seq libraries were sequenced on an Illumina HiSeq X Ten platform, and reads were aligned to the human or mouse genome using TopHat (v2.0.14). The expression levels were calculated using the Cufflinks software (v2.2.1) with default parameters. All RNA-seq experiments were performed with at least two biological replicates.

### Screening single-cell CRISPR clones by DNA-fragment editing

The generation of the CRISPR single-cell clones by DNA-fragment editing was performed as previously described (45, 46). Briefly, HEC-1-B or HEK293T cells at ∼80% confluency were transfected with Cas9 plasmids (0.6 μg) and two *HS5-1*-NRSE-targeted sgRNA-expressing plasmids (1.2 μg) by Lipofectamine 2000 (Life Technologies) in a 12-well plate. Two days post transfection, puromycin (Sigma) was added at a final concentration of 2 μg/ml. Four days later, cells were changed to fresh medium without puromycin and cultured for another two days. The cells were then diluted and plated into 96-well plates with approximately one cell per well. Two weeks later, single-cell clones were marked manually under a microscope and screened for targeted deletion by PCR. At least two single-cell clones for each deletion were obtained. We screened for a total of 133 single-cell clones, and 4 homozygous clones were obtained and analyzed. Single-cell clones for each editing were confirmed by Sanger sequencing and the screening primers are shown in Supplementary Table S1

### Generation of HS5-1-NRSE-deleted mice by CRISPR DNA-fragment editing

The generation of CRISPR mice was performed as previously described (33, 45). Briefly, Cas9 mRNA and a pair of sgRNAs targeting the NRSE sequences were injected into zygotes. Cas9 mRNA for mouse zygote injection was *in-vitro* transcribed from the XbaI-linearized Cas9 plasmid with a T7 promoter using mMESSAGE mMACHINE T7 Ultra Kit (Life Technologies). The sgRNAs were *in-vitro* transcribed from PCR products with the MEGAshortscript Kit (Life Technologies). The PCR products were amplified with a forward primer containing a T7 promoter followed by targeting sequences and a common reverse primer. Then, the transcribed RNA was purified with the MEGAclear Kit (Life Technologies) and eluted with the TE buffer (0.2 mM EDTA).

Zygotes obtained from the oviducts of super-ovulated C57BL/6 mice were injected with Cas9 mRNAs (100 ng/μl) and a pair of sgRNAs (50 ng/μl each). After culturing with KSOM medium (Millipore) for 0.5h at 37 °C in a 5% CO_2_ incubator, the injected embryos were transplanted into the oviducts of pseudo-pregnant ICR foster mothers. The F0 mice with *HS5-1* NRSE deletion were maintained and crossed to obtain F1 mice. F1 mice were genotyped again for heterozygous deletion. Heterozygous mice were then crossed to obtain homozygous mice. The wildtype littermates were used as controls.

For genotyping, mouse tails were lysed with 40 μl of the alkaline lysis buffer (25 mM NaOH and 0.2 mM disodium EDTA, pH12.0) at 98 °C for 40 min, and then neutralized with equal volume of the neutralizing buffer (40 mM Tris–HCl, pH5.0). 1 μl of the solution containing genomic DNA was used as template for PCR in a total volume of 20 μl to screen for NRSE deletion with specific primers (Supplementary Table S1).

### ChIP-seq

ChIP was performed as previously described (31, 33) with some modifications. Briefly, The P0 mouse tissues were dispersed by 0.0125% (w/v) (for brain) or 0.0625% (w/v) (for kidney) collagenase (Sigma) treatment in DMEM supplemented with 10% (v/v) FBS for 20 min at 37 °C with rotating at 700 rpm. Cells were then filtered through a 100 μm cell strainer (CORNING) to obtain single-cell suspension.

About 5 x 10^6^ cells were cross-linked with 1% formaldehyde for 10 min at room temperature, followed by quenching the crosslinking with glycine at a final concentration of 0.125 M and then spun down. Subsequently, cell pellets were lysed twice with the ChIP buffer (10 mM Tris-HCl, pH 7.5, 0.15 M NaCl, 1% Triton X-100, 1 mM EDTA, 0.1% SDS, 0.1% sodium deoxycholate, 1x protease inhibitors) by incubating at 4 °C for 10 min with slow rotation. Nuclei were spun down and resuspended in 400 μl of the ChIP buffer (for human cells) or ChIP buffer with 0.4% SDS (for cells from mouse tissues), and then sonicated using Vibracell ultrasonic processor (Sonics) (25% maximum, a train of 20s on and 20s off for 15 cycles). The sonicated solution was diluted with the ChIP buffer (1:5), pre-cleared with 40 μl Protein A agarose beads (Millipore), and then immunoprecipitated with specific antibodies against REST/NRSF (Millipore, 4 μg for each reaction), CTCF (Millipore, 2.5 μg for each reaction), H3K4me3 (Millipore, 2.5 μg for each reaction), and H3K27ac (Abcam, 2.5 μg for each reaction). The protein–DNA complexes were enriched with 40 μl Protein A/G magnetic beads (Thermo) by incubating at 4 °C for another 3h with slow rotation, and washed once with 1 ml of ChIP buffer, once with 1 ml of ChIP buffer with 0.4M NaCl, once with ChIP buffer without NaCl, once with ml of LiCl buffer (250 mM LiCl, 5% NP-40, 0.5% Sodium deoxycholate), by incubating at 4 °C for 10 min with slow rotation. Then, the samples were eluted with elution buffer (50 mM Tris-HCl, pH 8.0, 10 mM EDTA, 1% SDS) at 65 °C, shaken at 1000 rpm for 30 min, and then reverse-crosslinked by heating at 65 °C overnight. The DNA was purified with phenol-chloroform and precipitated with ethanol, and then used for library preparation.

Libraries were prepared according to the manual of Universal DNA Library Prep Kit for Illumina V2 (Vazyme). Briefly, DNA was end-repaired using End Prep Mix 3, ligated with index-containing adaptors using rapid DNA ligase in the rapid ligation buffer (66 mM Tris-HCl pH7.6, 20 mM MgCl2, 2 mM DTT, 2 mM ATP, 7.5% PEG 6000), amplified using primers and amplification mix to generate libraries. The library fragments of ∼350 bp (insert plus adaptor and PCR primer sequences) were selected and isolated with Agencourt AMPure XP beads (Beckman). Library DNA was sequenced on an Illumina HiSeq X Ten platform. The reads were sorted by indexes, aligned to hg19 or mm9 using Bowtie2, and peak-called using MACS2. All of the ChIP-seq experiments were performed with at least two biological replicates.

### Quantitative high-resolution chromosome conformation capture copy (QHR-4C)

QHR-4C experiments were performed as previously described (33) with some modifications. Briefly, cells were harvested from mice as described in ChIP-seq experiments. Then, about 2 million cells were crosslinked with 1% formaldehyde for 10 min at room temperature and the crosslinking reaction was quenched by glycine at a final concentration of 0.125 M. After spun down and washed twice with 1 x PBS, cell pellets were lysed twice with cold lysis buffer (50 mM Tris-HCl, pH 7.5, 150 mM NaCl, 1% Triton X-100, 5 mM EDTA, 0.5% NP-40, 1 x protease inhibitors). The pellet was resuspended in 225 μl of water, 30 μl of 10x DpnII buffer, and 7.5 μl of 10% SDS and incubated at 37 °C for 1h with constant shaking at 900 rpm. After quenching by adding 37.5 μl of 20% Triton X-100 and incubating at 37 °C for 1h with shaking at 900 rpm, the nuclei were then digested with 10 μl of DpnII (10 unit/μl) *in situ* at 37 °C with shaking at 900 rpm overnight. After enzyme inactivation at 65 °C for 20 min, nuclei were centrifuged at 2500 g for 5 min, resuspended and ligated in 358 μl of water, 40 μl of 10x T4 ligation buffer, and 2 μl of T4 DNA ligase (400 unit/μl) overnight at 16 °C. The ligated product was then reverse crosslinked by adding 10 μl of proteinase K (10 mg/ml) and heating at 65 °C for 4h, followed by adding 2 μl of RNase A (NEB) and incubating at 37 °C for 45 min. DNA was then extracted using phenol-chloroform. 2 μl of glycogen (20 mg/ml) was added to facilitate DNA precipitation. The precipitated DNA was dissolved in 50 μl water, and then sonicated to 200∼600 bp using the Bioruptor system with low energy setting at a train of 30s on with 30s off for 12 cycles.

With the fragmented DNA as template, a linearized amplification step was applied using a 5’ biotin-tagged primer (Supplementary Table S1) complementary to the viewpoint fragment in 100 μl of PCR system. The PCR product was denatured at 95 °C for 5 min and immediately chilled on ice to obtain ssDNA. The ssDNA was enriched and purified with Streptavidin Magnetic Beads (Thermo) according to the manufacturer’s instructions, and then ligated with annealed adaptors in 45 μl of the ligation system (20 μl of DNA-coated beads, 4.5 μl of 10x T4 ligation buffer, 10 μl of 30% PEG 8000, 1 μl of 50 μΜ adaptor, 0.9 μl of T4 DNA ligase, 8.6 μl of water). The beads were then washed twice to remove the free adaptors with the B/W buffer (5 mM Tris-HCl, 1M NaCl, 0.5 mM EDTA, pH7.5). The DNA-coated beads were resuspended in 10 μl water. With the DNA on beads as a template, the QHR-4C libraries were generated by PCR amplification with a pair of primers (Supplementary Table S1). Finally, the libraries were purified with a PCR purification kit (Qiagen) and sequenced on an Illumina HiSeq X Ten platform. Reads were sorted using indexes and barcodes, mapped to the mouse (NCBI37/mm9) or human (GRCh37/hg19) genomes using Bowtie2, and calculated using the r3Cseq program (version 1.20) in the R package (version 3.3.3). All of the QHR-4C experiments were performed with at least two biological replicates.

### Structural modeling and molecular dynamics (MD) simulations

Structural modeling and MD simulations were performed as previously described (55). Briefly, we first built the models of REST-*HS5-1*-NRSE and REST-*PCDHγa6*-NRSE based on one crystal structure of the ZF protein PR/SET domain 9 (PRDM9) in complex with DNA (PDB id: 5v3g). The REST/NRSF structure was created using the homology modeling strategy and DNA was modeled by directly replacing the DNA nucleobases in 5v3g to the ones of *HS5-1*-NRSE or *PCDHγa6*-NRSE. In particular, the orientation of the REST-ZF6 in REST-*HS5-1*-NRSE in respect to DNA was constructed according to the formerly captured CTCF-ZF8 structure that displays a non-DNA-penetration conformation (56, 57).

Then, the above two DNA-bound REST/NRSF complexes were subject to energy minimization, followed by MD simulations. The protein/DNA were described using the AMBER force field ff14SB, with the bsc1 corrections used for the DNA nucleotides. Each solute was immersed in a cubic box filled with the TIP3P waters, and appropriate number of Na^+^ ions were added to neutralize the whole system. The final structure includes a total of 49,652 and 68,845 atoms for REST-*PCDHγa6*-NRSE and REST-*HS5-1*-NRSE, respectively. Energy minimization was conducted using the steepest decent and conjugate gradient methods. Then, each complex was heated from 0 to 310 K within 200-ps MD simulations, followed by 200-ps equilibrium MD simulations by constraining all the solute heavy-atoms. Finally, we performed three parallel 50-ns production MD simulations, each initiated from different velocities. The temperature was controlled by the Langevin thermostat at 310 K and the SHAKE algorithm was used to constrain the bond lengths involving hydrogen atoms. The non-bonded cutoff distance was set as 12 Å, and the long-range electrostatic interactions were treated by the particle mesh Ewald (PME) method. All the MD simulations were conducted using the AMBER 2018 package.

## SUPPLEMENTARY DATA

Supplementary Data are available.

## ACKNOWLEDGMENTS

We thank all members of our laboratory for helpful discussions and critical reading of the manuscript.

## FUNDING

This study was supported by grants to Q.W. from the National Natural Science Foundation of China (31630039 and 91940303), the Ministry of Science and Technology of China (2017YFA0504203 and 2018YFC1004504), and the Science and Technology Commission of Shanghai Municipality (19JC1412500).

## CONFLICT OF INTEREST

The authors declare no competing interests.

## SUPPLEMENTARY FIGURES LEGENDS

**Supplementary Figure S1.**
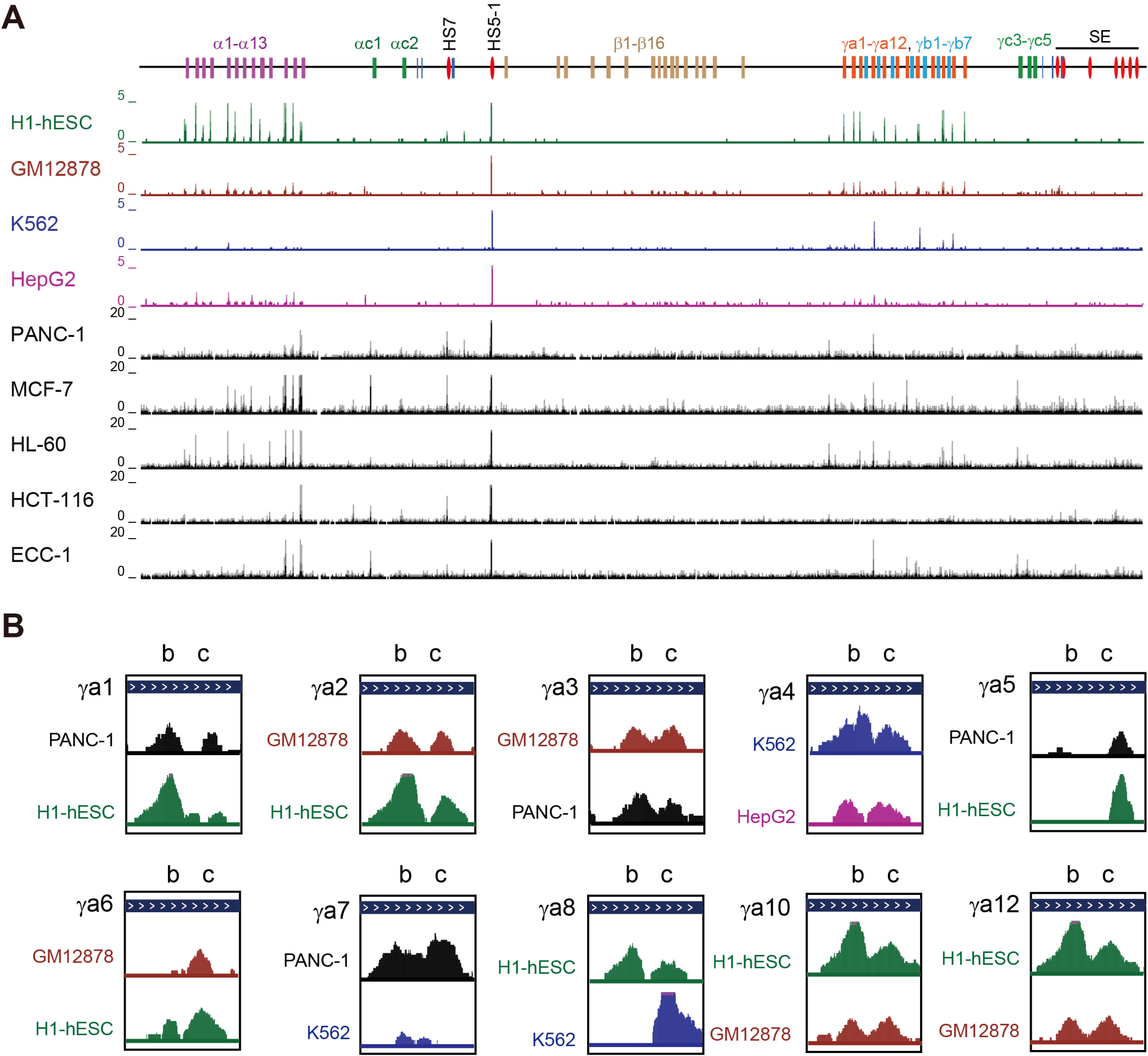
REST/NRSF binding patterns in the three human PCDH clusters. (**A**) REST/NRSF binding in clustered *PCDH* in different cell lines (data are from the ENCODE project). (**B**) Showing are the two REST/NRSF peaks in the *PCDHγa* alternate exons.

**Supplementary Figure S2.**
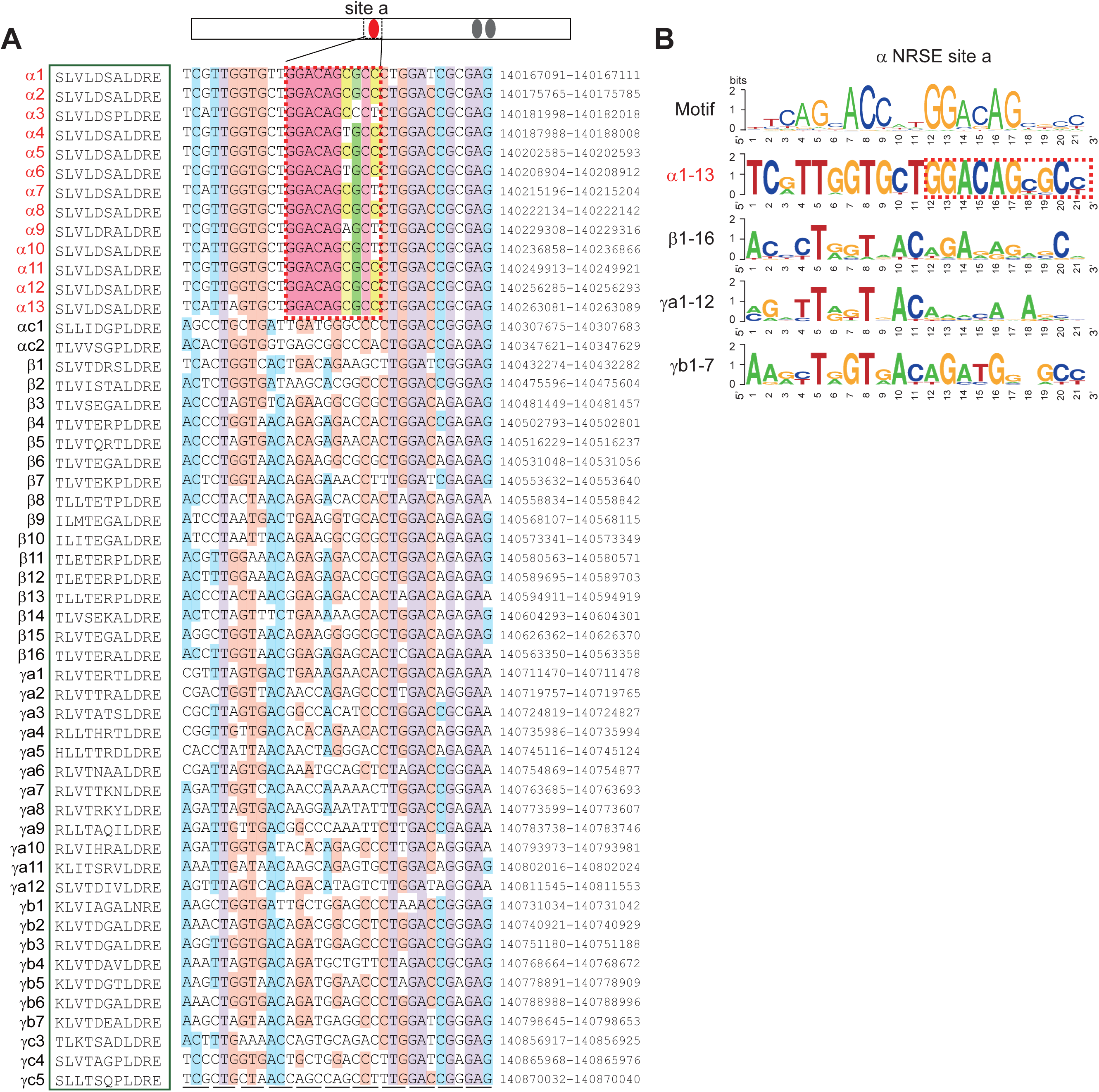
Sequence alignment of site ‘a’ in clustered *PCDH* variable exons. (**A**) Alignment and genomic location of the sequences in the ‘a’ site of the clustered *PCDH* variable exons. (**B**) Comparison of site ‘a’ consensus sequences of *PCDH* subclusters. Note that only alternate members of the *PCDHα* cluster contains NRSE site ‘a’.

**Supplementary Figure S3.**
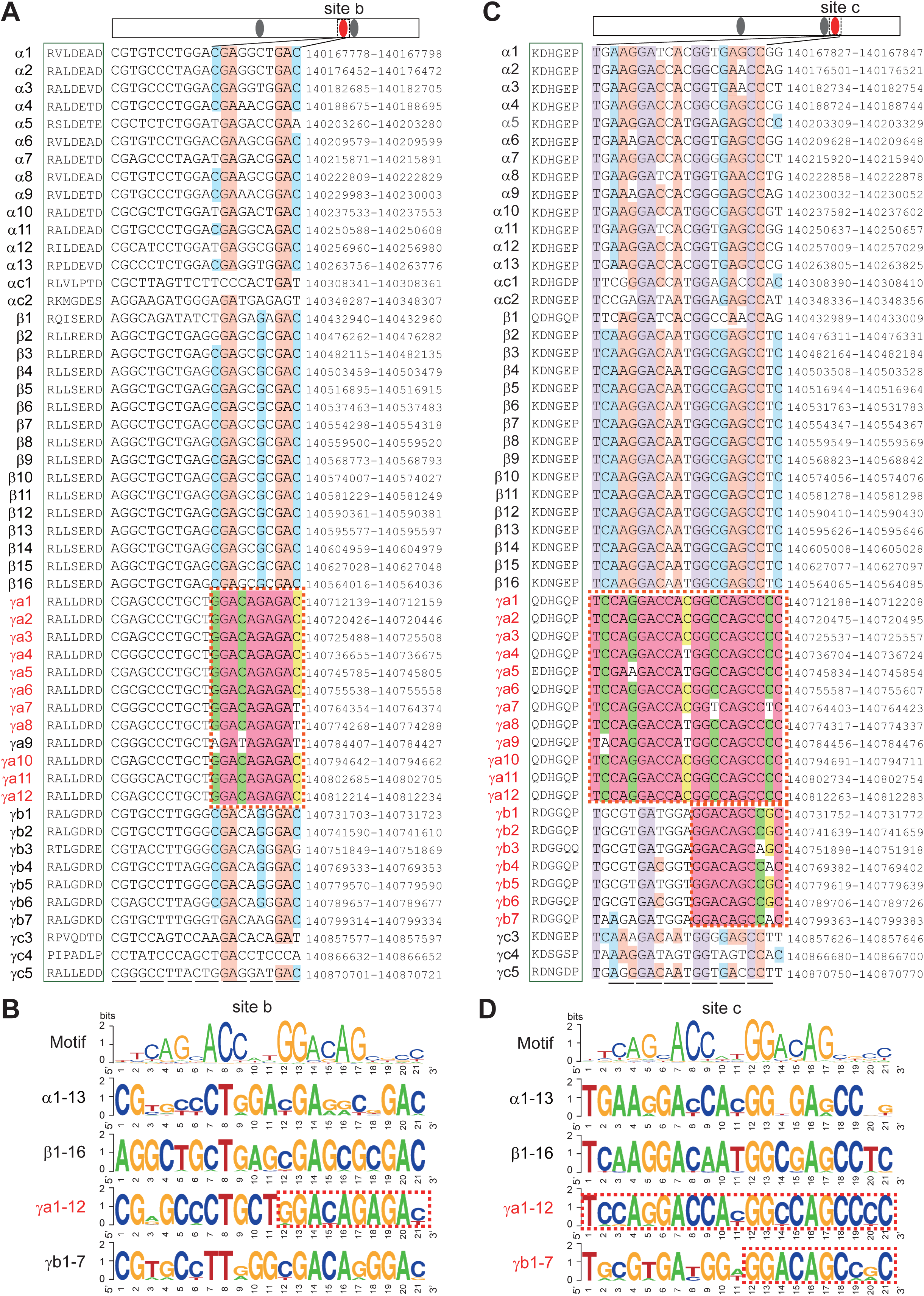
Sequence alignment of sites ‘b’ and ‘c’ in clustered *PCDH* variable exons. (**A**) Alignment and genomic location of the sequences in the site ‘b’ of the clustered *PCDH* variable exons. (**B**) Comparison of site ‘b’ consensus sequences of *PCDH* subclusters. (**C**) Alignment and genomic location of the sequences in the site ‘c’ of the clustered *PCDH* variable exons. (**D**) Comparison of site ‘c’ consensus sequences of *PCDH* subclusters.

**Supplementary Figure S4.**
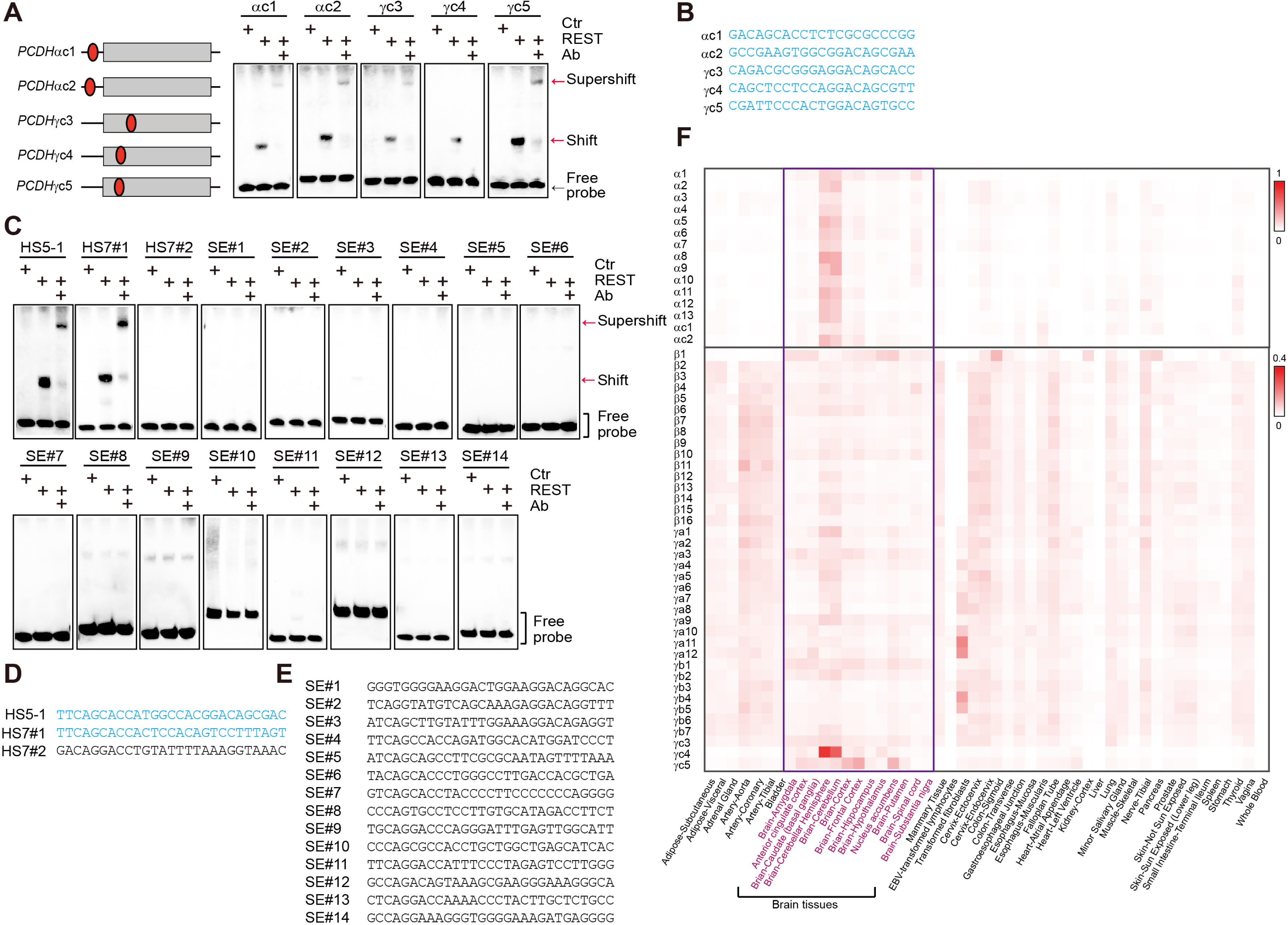
REST/NRSF binding profiles in C-type genes and super-enhancers. (**A**) EMSA assays of REST/NRSF with the NRSE probes of C-type *PCDH* genes, using mock protein as a control. (**B**) The sequences of the NRSE motif in the C-type *PCDH* genes. (**C**) EMSA assays of REST/NRSF with putative NRSE probes within the super-enhancers, using mock protein as a control. (**D**) Probe sequences in the super-enhancer of *PCDHα*. (**E**) Probe sequences in the super-enhancer of *PCDHβγ*. (**F**) The expression levels of the clustered *PCDH* genes in different tissues. (41)

**Supplementary Figure S5.**
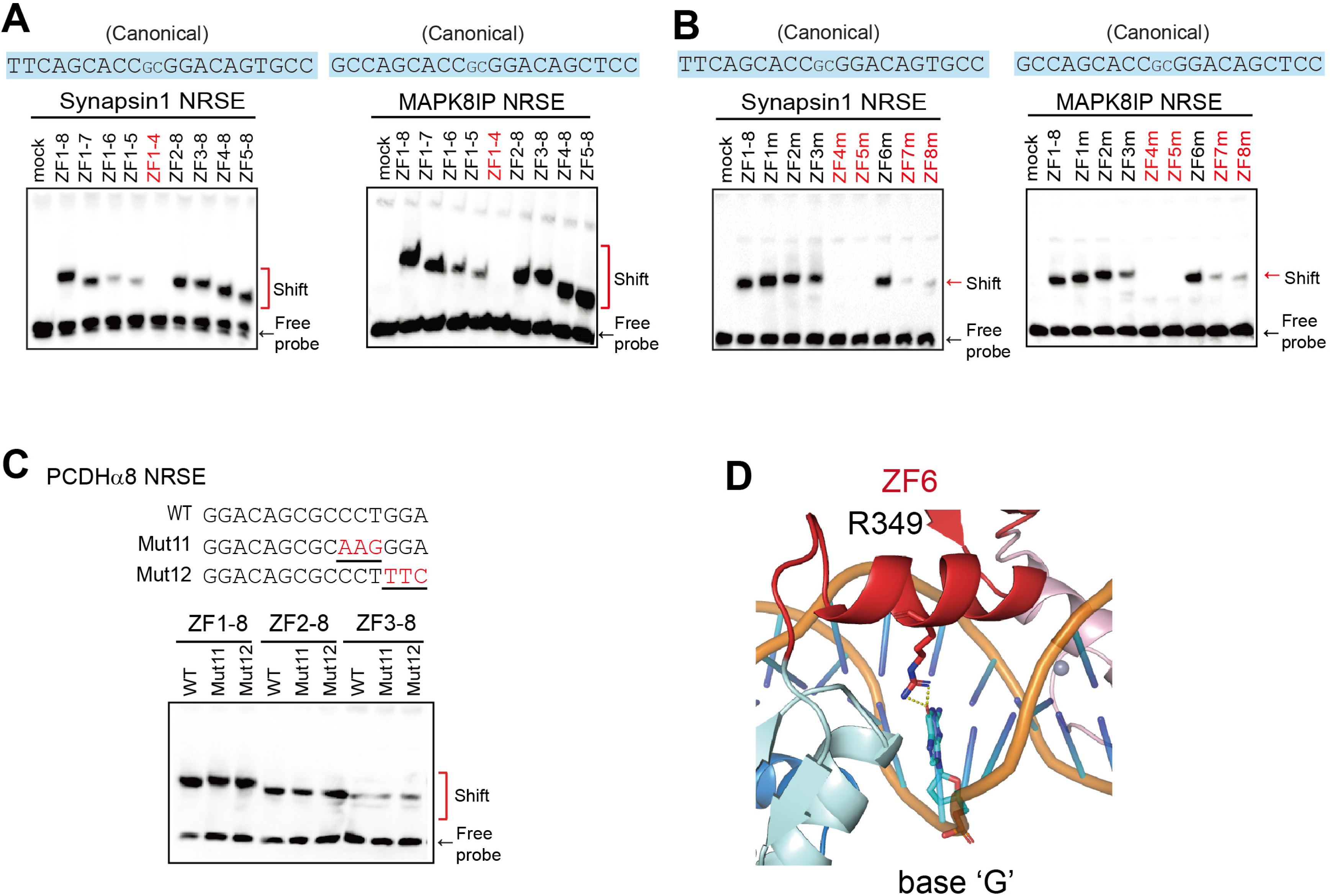
Binding patterns of ZF-mutated or ZF-deleted REST/NRSF with different probes. (**A**) EMSA assays for the binding of ZF-deleted REST/NRSF with two canonical NRSEs. (**B**) EMSA assays for the binding of ZF-mutated REST/NRSF with two canonical NRSEs. (**C**) EMSA assays for the ZF-deleted REST/NRSF with the Mut11 and Mut12 probes of *PCDHα8* NRSE. (**D**) Shown are the contacts of R349 in ZF6 with the base ‘G’ in the complementary strand at the 9^th^ position of the canonical *PCDHγa6* NRSE.

**Supplementary Figure S6.**
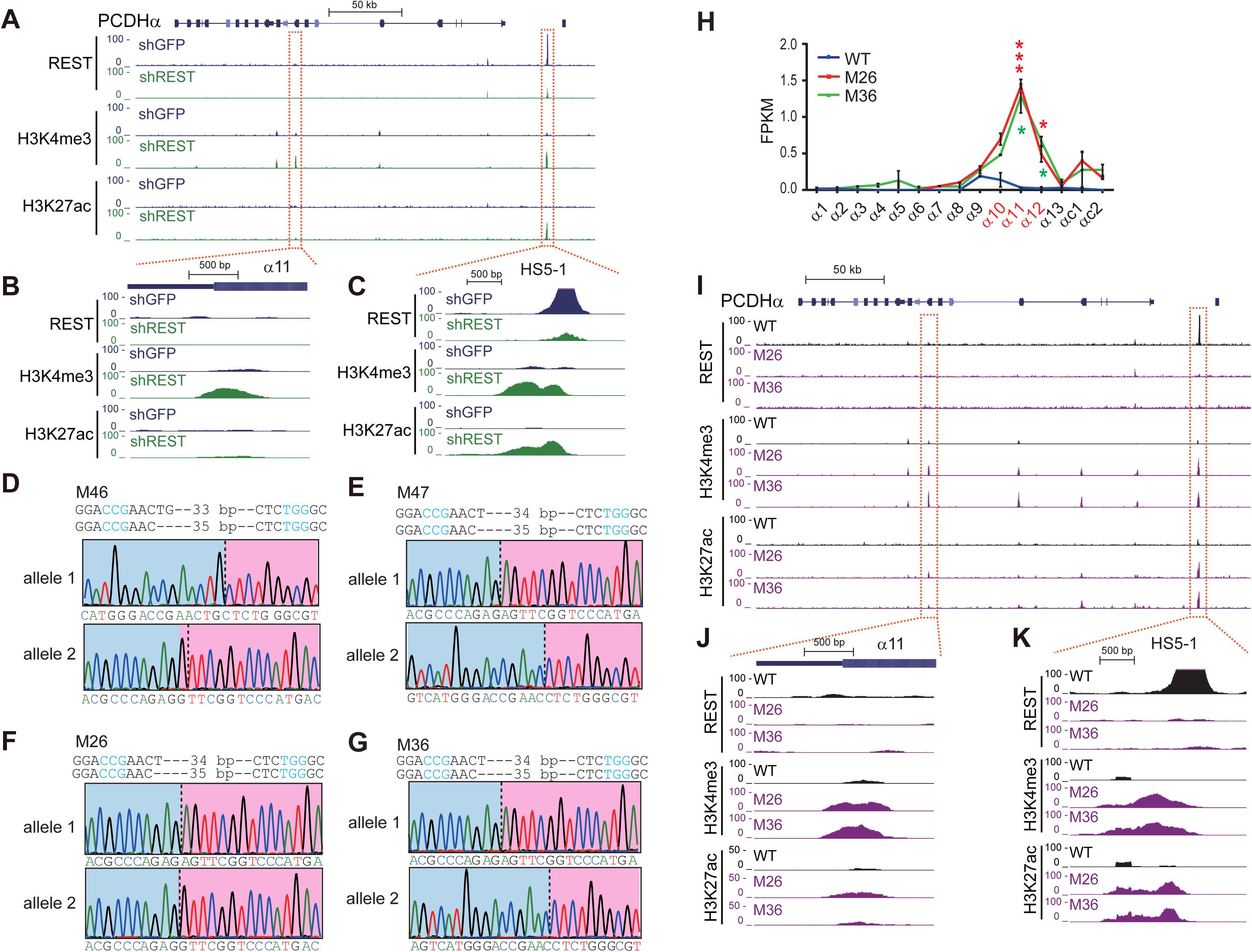
Genotyping and CRISPR screening of single-cell NRSE deletion clones by DNA-fragment editing in HEK293T and HEC-1-B cells. (**A**-**C**) ChIP-seq with a specific antibody against REST/NRSF, H3K4me3, or H3K27ac in HEK293T cells upon knockdown of REST/NRSF by shRNA. (**D**, **E**) Genotyping of the *HS5-1* NRSE-deleted single-cell clones of HEC-1-B cells (M46, M47). The PAM sites are highlighted. (**F**, **G**) Genotyping of the *HS5-1* NRSE-deleted single-cell clones of HEK293T cells (M26, M36). The PAM sites are highlighted. (**H**) Expression profiles of *PCDHα* measured by RNA-seq in HEK293T single-cell CRISPR clones. Data are presented as mean ± SEM. *\* P < 0.05*, *\*\*\* P < 0.001*. (**I**-**K**) ChIP-seq of REST/NRSF, H3K4me3, and H3K27ac in HEK293T single-cell CRISPR clones upon deletion of the *HS5-1* NRSE.

**Supplementary Figure S7.**
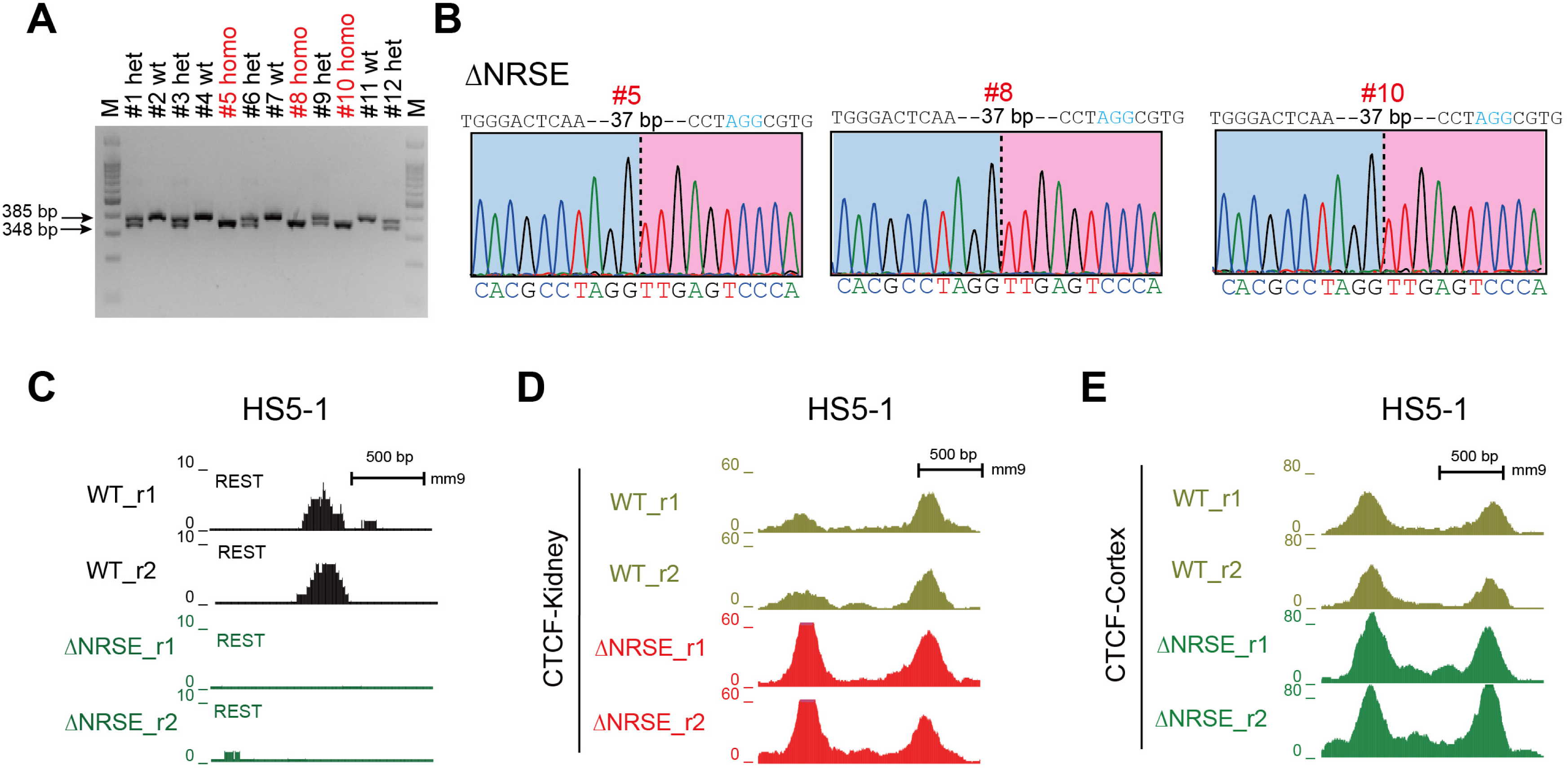
Genotyping and CTCF binding of the *HS5-1* NRSE-deleted mice by DNA-fragment editing. (**A**, **B**) Genotyping of P0 mice with the *HS5-1* NRSE deletion. (**C**) ChIP-seq confirmed the abolishment of REST/NRSF binding in the *HS5-1* enhancer upon NRSE deletion in mice. (**D**) CTCF ChIP-seq peak of *HS5-1* in the kidney tissues from WT and *HS5-1* NRSE-deleted mice. (**E**) CTCF ChIP-seq peak of *HS5-1* in the cortical tissues from WT and *HS5-1* NRSE-deleted mice.

Supplementary Table S1. List of Primers Used.

## Notes

### Competing Interest Statement

The authors have declared no competing interest.

