## Supplemental Table 1 for "Mechanism of REST/NRSF Regulation of Clustered Protocadherin α Genes"

**Supplementary Table S1. List of Primers Used**

| Name | 5'-3' sequence |
|---|---|
| Primers used to generate plasmids for *in-vitro* expression of REST/NRSF proteins | |
| hREST(ZF1-8)-F | CGGAATTCACCATGGGTGCTCCAGATATTTACA |
| hREST(ZF2-8)-F | CGGAATTCACCATGTCCAAGGGCCCCATTCGC |
| hREST(ZF3-8)-F | CGGAATTCACCATGAATGAGCGAGTCTACAAG |
| hREST(ZF4-8)-F | CGGAATTCACCATGCCAAGGAAAGTATACACA |
| hREST(ZF5-8)-F | CGGAATTCACCATGGGAGAACGCCCATATAAAT |
| hREST(ZF1-8)-R | GCTCTAGATTACAGATCCTCTTCTGAGATGAGTTTTTG<br>TTCGG TAATATTATCAGGCAA |
| hREST(ZF1-7)-R | GCTCTAGATTACAGATCCTCTTCTGAGATGAGTTTTTG<br>TTCATTGAACTGCCGTGGGTT |
| hREST(ZF1-6)-R | GCTCTAGATTACAGATCCTCTTCTGAGATGAGTTTTTG<br>TTCAT<br>TAAGAGGTTTAGGCCC |
| hREST(ZF1-5)-R | GCTCTAGATTACAGATCCTCTTCTGAGATGAGTTTTTG<br>TTCTTTAAATGGCTTCTCACC |
| hREST(ZF1-4)-R | GCTCTAGATTACAGATCCTCTTCTGAGATGAGTTTTTG<br>TTCTTTATATGGGCGTTCTCC |
| hREST-ZF1m-F | ACCCTTTCGCCGTAAGCCACGCCAATATGA |
| hREST-ZF2m-R | TCATATTGGCGTGGCTTACGGCGAAAGGGT |
| hREST-ZF2m-F | CCCCATTCGCCGTGACCGCCGCGGCTACAA |
| hREST-ZF2m-R | TTGTAGCCGCGGCGGTCACGGCGAATGGGG |
| hREST-ZF3m-F | AGTCTACAAGCGTATCATTCGCACATACAC |
| hREST-ZF3m-R | GTGTATGTGCGAATGATACGCTTGTAGACT |
| hREST-ZF4m-F | AGTATACACACGCGGAAAACGCAACTATTT |
| hREST-ZF4m-R | AAATAGTTGCGTTTTCCGCGTGTGTATACT |
| hREST-ZF5m-F | CCCATATAAACGGGAACTTCGTCCTTACTC |
| hREST-ZF5m-R | GAGTAAGGACGAAGTTCCCGTTTATATGGG |
| hREST-ZF6m-F | GCCATTTAAACGCGATCAGCGCAGTTATGT |
| hREST-ZF6m-R | ACATAACTGCGCTGATCGCGTTTAAATGGC |
| hREST-ZF7m-F | ACCTCTTAATCGCCCACACCGTGATTACAA |
| hREST-ZF7m-R | TTGTAATCACGGTGTGGGCGATTAAGAGGT |
| hREST-ZF8m-F | GCAGTTCAATCGCCCTGTACGTGACTATGC |
| hREST-ZF8m-R | GCATAGTCACGTACAGGGCGATTGAACTGC |
| Primers used to clone the wild type REST/NRSF binding sites for EMSA | |
| EMSA-hPCDHα1-F | AGAACTGGCGGTCACTTCAT |
| EMSA-hPCDHα1-R | AGCTCACTGACTGCACCAATAG |
| EMSA-hPCDHα2-F | CCATCTCAGAGAACGCTTCC |
| EMSA-hPCDHα2-R | CTGCCAGTACCCAGCTGAAG |
| EMSA-hPCDHα3-F | GATCACTGCACAGTTCTACTCG |
| EMSA-hPCDHα3-R | GAGGAAGGCTAGGGCTAAAAG |
| EMSA-hPCDHα4-F | GCGTGTCCGACAAAGACATG |

| | |
|---|---|
| EMSA-hPCDHα4-R | CCTGTCCACTCTCCACAAGT |
| EMSA-hPCDHα5-F | GCCATAACCACCCTTTTCC |
| EMSA-hPCDHα5-R | TTGTAGCCCGAATCAGGGT |
| EMSA-hPCDHα6-F | TAATAGCCTTGTTGCAGCC |
| EMSA-hPCDHα6-R | GTGATGACGCCTTTGGAG |
| EMSA-hPCDHα7-F | CCAGGTACCGTCATCACATTG |
| EMSA-hPCDHα7-R | GTCAGCGTCAACTGCACGT |
| EMSA-hPCDHα8-F | TCAGAATCCAGAATGCCAGAC |
| EMSA-hPCDHα8-R | CTCATAGGCCGACACTCTC |
| EMSA-hPCDHα9-F | TTAGTGTGATCGACCTAGACG |
| EMSA-hPCDHα9-R | ACGCCCGCGACGATGACTTT |
| EMSA-hPCDHα10-F | CCTAATCAGCGTTTCTGACCAT |
| EMSA-hPCDHα10-R | CACATCTACCATTGGGCATG |
| EMSA-hPCDHα11-F | TCTCCTGAAGTCGCCGTG |
| EMSA-hPCDHα11-R | GACCGCGGTACTAGCTTGTT |
| EMSA-hPCDHα12-F | CAATGTCCCTGAAGTAATGGTTAC |
| EMSA-hPCDHα12-R | CGTAGGACAGCCAAGCGTTAT |
| EMSA-hPCDHα13-F | TAACGCCCCAGAGGTTACC |
| EMSA-hPCDHα13-R | CCCTCGACGAAGCCTGT |
| EMSA-hPCDHβ4-F | ATGCTCCTGAGACGGTAGTCT |
| EMSA-hPCDHβ4-R | TCAGGAACTTGAACTCACCAG |
| EMSA-hPCDHβ7-F | ATCGACCCGAGCTGCTCC |
| EMSA-hPCDHβ7-R | GCCTGTGCTCTGGGGTAGC |
| EMSA-hPCDHγa1-F | CTCTTATCAGTGTGCATGACCAG |
| EMSA-hPCDHγa1-R | GTAAAGACTGGGGTGCTGATAG |
| EMSA-hPCDHγa2-F | GTTGCGAGACTTGCAAGTGT |
| EMSA-hPCDHγa2-R | TTCGAGTAGAGATTGAGGCG |
| EMSA-hPCDHγa3-F | CAACGCCCGCATCACTTATG |
| EMSA-hPCDHγa3-R | TGCGAGCCGGGCGTACT |
| EMSA-hPCDHγa4-F | CACTTTTCAACGTGCATGACAGT |
| EMSA-hPCDHγa4-R | TGAGGCCCGAGTCGTCTG |
| EMSA-hPCDHγa5-F | GCTGTTTAGCGTACATGATGGT |
| EMSA-hPCDHγa5-R | GGTCAATGGGGGTCTTGATAC |
| EMSA-hPCDHγa6-F | GGAATTCAACATGGCGCCTCCGCAGA |
| EMSA-hPCDHγa6-R | GGGAAGCTTTTACTTCTTCTCCTTCTTGCCCG |
| EMSA-hPCDHγa7-F | GTGACTGCACATGACAGCG |
| EMSA-hPCDHγa7-R | CACGGTGAGTGTGACGGT |
| EMSA-hPCDHγa8-F | CTGTGACCGAGGACACGCT |
| EMSA-hPCDHγa8-R | GTGTAAGGCTCGAATCGTTCG |
| EMSA-hPCDHγa9-F | GAGATTTGCAAATGCAGGTG |
| EMSA-hPCDHγa9-R | TACTGTGAGCGTGACAGTGG |
| EMSA-hPCDHγa10-F | CTTCTCAGTGACAGCGCTG |
| EMSA-hPCDHγa10-R | GGAACCAATGGGGTATCCTC |
| EMSA-hPCDHγa11-F | GATTGCTCTTCTAAATGTGCAAG |

| | |
|---|---|
| EMSA-hPCDHγa11-R | CACCACCAAGTACAGCGAG |
| EMSA-hPCDHγa12-F | CACCATCCAAGGGGCAAG |
| EMSA-hPCDHγa12-R | AATACCGAATCACCTGACAGC |
| EMSA-hPCDHγb1-F | GGGACCCAACGGCAGAG |
| EMSA-hPCDHγb1-R | AGCAGCCCTCAGTGTCGAG |
| EMSA-hPCDHγb2-F | CCTGACTTGGGCCCCAGT |
| EMSA-hPCDHγb2-R | GGGGTCAGAGGGCTCCC |
| EMSA-hPCDHγb3-F | CCTGGCTTCTGAATCCCAAC |
| EMSA-hPCDHγb3-R | GAAACTGTAGCTCCGCCTGAG |
| EMSA-hPCDHγb4-F | CTATTTCACAAGTCAGGGCTTC |
| EMSA-hPCDHγb4-R | AACCACGGATTCGGATTTAAC |
| EMSA-hPCDHγb5-F | GACCTAGAGCCTCTGGCAC |
| EMSA-hPCDHγb5-R | GATGGGAAGTCGACTCGC |
| EMSA-hPCDHγb6-F | GGGCTCAATGGCCACAT |
| EMSA-hPCDHγb6-R | GATGGAAGCAGTCCCAAGTAG |
| EMSA-hPCDHγb7-F | ACCTGGAGTCACGAACGC |
| EMSA-hPCDHγb7-R | GAGTCAGAGGGTGTGGGATG |
| EMSA-hPCDHαc1pro-F | TAAGATCTGGGCAGCCTCAG |
| EMSA-hPCDHαc1pro-R | GTCCACGTTCCACCAACAC |
| EMSA-hPCDHαc1-F | CCTCCCAGAAGTGCAACAG |
| EMSA-hPCDHαc1-R | CAGAAAGCTGCCCTCCTG |
| EMSA-hPCDHαc2pro-F | GGGAGCTGATAGCCAGACTTC |
| EMSA-hPCDHαc2pro-R | CGCTCACGCTCTAGAAAGC |
| EMSA-hPCDHγc3-F | GAGATTAGCGAGGCCGTG |
| EMSA-hPCDHγc3-R | CGGACACTTGAACACGCAC |
| EMSA-hPCDHγc4-F | GTGAACCAAAGACACTTCCGT |
| EMSA-hPCDHγc4-R | ACCAGGCGGTAGTCCGAT |
| EMSA-hPCDHγc5-F | CTATTTTTCCCTGAGCTTGATG |
| EMSA-hPCDHγc5-R | TCCCACACGTAGAACTGAGG |
| EMSA-hPCDHHS7-F | GGATTAACTCTTGGCAGTCCTG |
| EMSA-hPCDHHS7-R | GCAGAGCTGACACAATAGCAC |
| EMSA-hPCDHSE#1/2-F | CCGTATCACTGATACCTTGGC |
| EMSA-hPCDHSE#1/2-R | TGAGTGATGTATGCATGGGG |
| EMSA-hPCDHSE#3/4/5-F | GCTGCAACAGGACAAGATCC |
| EMSA-hPCDHSE#3/4/5-R | CCAAAGGTCAGGCATCAGG |
| EMSA-hPCDHSE#6/7/8-F | GGTGACCCCTATATTCCCAGTGC |
| EMSA-hPCDHSE#6/7/8-R | TTCTCTGGCAGCCCCGCT |
| EMSA-hPCDHSE#9/10-F | TGCTTGTGAAGTCCAGAGTAAGG |
| EMSA-hPCDHSE#9/10-R | CCGGGTCAAGCAAATGAG |
| EMSA-hPCDHSE#11/12/13-F | TGATGGACATAGGGGTGTTCAT |
| EMSA-hPCDHSE#11/12/13-R | TTTATCTTGGGCAGAGCAAGTAG |
| EMSA-hPCDHSE#14-F | CTGTATCTCACCCACGCTAACC |
| EMSA-hPCDHSE#14-R | GCAAGATGGGATGGAACCAT |
| EMSA-hHS5-1-F | TACCGCTCGAGAATACTACTAGGCACTTGTTTGGATC |

| | |
|---|---|
| EMSA-hHS5-1-R | GAGCGAGCTCTGGCTAACAACATAGTGCTTCCT |
| EMSA-CELSR3-F | TCGCCGAGGTTACTTTCCTG |
| EMSA-CELSR3-R | TTTTTGATTCGGCACCACGG |
| EMSA-MAPK8IP-F | ACATCCTCTCCTTTCTCTGCC |
| EMSA-MAPK8IP-R | GCAGCCAATGCGGATCAGT |
| EMSA-synapsin1-F | CCTAGATTGGCGTGTGTTCTG |
| EMSA-synapsin1-R | AGGTAGTTCATGGCTGCGAC |
| Primers used to mutate REST/NRSF binding sites for EMSA | |
| EMSA-hPCDHα8-M1R | TAGGCCGACACTCTCTCGCGGTCCAGGGCGCTGGAAAGCAC |
| EMSA-hPCDHα8-M2R | CTCATAGGCCGACACTCTCTCGCGGTCCAGGGCGAGTTCC |
| EMSA-hPCDHα8-M3R | TAGGCCGACACTCTCTCGCGGTCCAGGTATCTGTCCAGCAC |
| EMSA-hPCDHα8-M11R | TAGGCCGACACTCTCTCGCGGTCCCTTGCGCTGTCC |
| EMSA-hPCDHα8-M12R | TAGGCCGACACTCTCTCGCGGGAAAGGGCGCTGTCC |
| EMSA-hHS5-1-M1R | TAAAGGAAATCTCTCTGGCCACGCCCAGAGTTCAGCACCATGGCCACTTCCAGCG |
| EMSA-hHS5-1-M2R | TAAAGGAAATCTCTCTGGCCACGCCCAGAGTTCAGCACCATGGCCACGGAACTCGACA |
| EMSA-hHS5-1-M3R | TAAAGGAAATCTCTCTGGCCACGCCCAGAGTTCAGCACCATGGCCACGGACAGATCCAC |
| EMSA-hHS5-1-M4R | TAAAGGAAATCTCTCTGGCCACGCCCAGAGTTCAGCCAAATGGC |
| EMSA-hHS5-1-M5R | TAAAGGAAATCTCTCTGGCCACGCCCAGAGTTCCTAACCAT |
| EMSA-hHS5-1-M6R | TAAAGGAAATCTCTCTGGCCACGCCCAGAGGGAAGCAC |
| EMSA-hHS5-1-M7R | TAAAGGAAATCTCTCTGGCCACGCCCAGAGTTCAGCACACGGGCCA |
| EMSA-hHS5-1-M8R | TAAAGGAAATCTCTCTGGCCACGCCCAGAGTTCAGACACATGG |
| EMSA-hHS5-1-M9R | TAAAGGAAATCTCTCTGGCCACGCCCAGAGTTACTCACCA |
| EMSA-hHS5-1-M10R | TAAAGGAAATCTCTCTGGCCACGCCCAGATGGCAGCA |
| Primers used to amplify probes for EMSA | |
| hPCDHα1/2/4/5/6/8/11a-NRSE-F† | CACCTTCAAGAATTACTACTCG |
| hPCDHα1/5a-NRSE-R | CTCATAGACCGACAGGCT |
| hPCDHα2/4a-NRSE-R | CTCATAGGCTGACACGCT |
| hPCDHα3/6/12a-NRSE-R | CTCATAGGCCGACACGCT |
| hPCDHα8a-NRSE-R | CTCATAGGCCGACACTCTC |
| hPCDHα11a-NRSE-R | TTCATAGGCCCACACGTT |
| hPCDHα3/10a-NRSE-F† | CACCTACAAGAATTACTA |
| hPCDHα10a-NRSE-R | CTCATAGGCCGACACCCT |
| hPCDHα7a-NRSE-F† | CACCTTCAAGAATTACTATTC |

| | |
|---|---|
| hPCDHα7a-NRSE-R | CTCATAGGCGGACACACT |
| hPCDHα9/12a-NRSE-F† | CACCTACAAGAATTACTACTCG |
| hPCDHα9a-NRSE-R | CTCGTAGGCGGACACACT |
| hPCDHα13a-NRSE-F† | CACCTACAAGAACTACTACTC |
| hPCDHα13a-NRSE-R | TTCATAGGCTGATACGCT |
| hPCDHβ7aCtr-NRSE-F† | CTTCTATACTCTGGTAACAG |
| hPCDHβ7aCtr-NRSE-R | GGTGATGGTGATGTTGTACT |
| hPCDHγa4aCtr-NRSE-F† | TTATTATCGGTTGTTGACAC |
| hPCDHγa4aCtr-NRSE-R | AGTTACAGTGATGTTATATTC |
| hPCDHγb3aCtr-NRSE-F† | CACATACAGGTTGGTGAC |
| hPCDHγb3aCtr-NRSE-R | TGTGATCGTCACATTGTATTC |
| hPCDHγa1/2/3/6/7/8/10/12b-NRSE-F† | GGGCGAGGTGCGCACGGC |
| hPCDHγa1b-NRSE-R | GACGGCCACCACGAGACT |
| hPCDHγa2b-NRSE-R | GATGGCCACCACGAGGCT |
| hPCDHγa3/5/7/8/11b-NRSE-R | GACGGCCACCACGAGGCT |
| hPCDHγa6b-NRSE-R | GACGGCCACCACTAGGCTCTGC |
| hPCDHγa10/12b-NRSE-R | GACGGCCACTACGAGGCT |
| hPCDHγa4b-NRSE-F† | AGGCGAGGTGCGCACCGC |
| hPCDHγa4b-NRSE-R | GACGACCACTACAAGCCT |
| hPCDHγa5b-NRSE-F† | GGGCGAGGTGCGCACAGC |
| hPCDHγa9b-NRSE-F† | AGGTGAAGTGCGCACAGC |
| hPCDHγa9b-NRSE-R | TACAGCCACCACAAGGCT |
| hPCDHγa11b-NRSE-F† | GGGCGAGGTGCGTACAGC |
| hPCDHβ4bCtr-NRSE-F† | TGGCGAGGTGCGCACCGC |
| hPCDHβ4bCtr-NRSE-R | GACAAGCACCACGAGCCT |
| hPCDHγb3bCtr-NRSE-F† | GGGTGAGGTGCGCACGGC |
| hPCDHγb3bCtr-NRSE-R | CACGGCGACCAGAAGGCG |
| hPCDHαc1bCtr-NRSE-F† | AGGTGAGCTCCGTACTGC |
| hPCDHαc1bCtr-NRSE-R | AACCACTACCACCACCCT |
| hPCDHγa1c-NRSE-F† | ACGCGCTCAAGCAGAGGC |
| hPCDHγa1/2/3/6/10/12c-NRSE-R | CCACGGTGAGCGTGACAGT |
| hPCDHγa2/3/5/6/11/12c-NRSE-F† | ACGCGCTCAAGCAGAGCC |
| hPCDHγa4c-NRSE-F† | ACGCGCTCAAGCAGAGGC |
| hPCDHγa4c-NRSE-R | CCACAGTGAGTGTGACGG |
| hPCDHγa5c-NRSE-R | CAACGGTGACCGTGAAGG |
| hPCDHγa7c-NRSE-F† | ATGCCCTCAAGCAGAGCC |
| hPCDHγa7c-NRSE-R | CCACGGTGAGTGTGACGGT |
| hPCDHγa8c-NRSE-F† | ATGCGCTCAAGCAGAGCC |
| hPCDHγa8c-NRSE-R | CTACGGTGAGCGTGACAG |
| hPCDHγa9c-NRSE-F† | ATGCGCTCAAACAGAGCC |
| hPCDHγa9c-NRSE-R | CTACTGTGAGCGTGACAG |
| hPCDHγa10c-NRSE-F† | ACGCGCTCAAGCAAAGCC |
| hPCDHγa11c-NRSE-R | CCACGGTGAGCGTGACGG |
| hPCDHγb1/2/5c-NRSE-F† | ACGCGGCCCGCCAGCGCC |

| | |
|---|---|
| hPCDHγb1c-NRSE-R | TTAGGTGCAGCGTGGCGG |
| hPCDHγb2c-NRSE-R | TTAGGTGCAGCGTGGCCG |
| hPCDHγb3c-NRSE-F† | AGGCCGCCCGCCAGCGCC |
| hPCDHγb3c-NRSE-R | TTAGGTGCAGCATGACGG |
| hPCDHγb4c-NRSE-F† | ACGCCGTCCGCCAGCGCC |
| hPCDHγb4c-NRSE-R | CCAGGTGCAACGTGGCAG |
| hPCDHγb5c-NRSE-R | CCAAGTGCAGCGTGGCGG |
| hPCDHγb6c-NRSE-F† | ACGCAGCCCGCCAGCGCC |
| hPCDHγb6c-NRSE-R | CCAGATGAAGCGTGGCGG |
| hPCDHγb7c-NRSE-F† | ACTCGGTCCGCCAGCGCC |
| hPCDHγb7c-NRSE-R | CCAGGTGCAGCGTGGCAG |
| hPCDHα4cCtr-NRSE-F† | ACGCTCCGCGCCACCGCC |
| hPCDHα4cCtr-NRSE-R | ACACCAGCACAGTGGCCG |
| hPCDHβ4cCtr-NRSE-F† | ACGCAGCCAAGCACAGGC |
| hPCDHβ4cCtr-NRSE-R | GCACGTGCAGCGTGGCGG |
| hPCDHαc1pro-NRSE-F† | TGGTCGAGACCCCAGCCC |
| hPCDHαc1pro-NRSE-R | CAGAGACGAGGCCGCCCG |
| hPCDHαc2pro-NRSE-F† | GATGGGGCTGGAGAGGCT |
| hPCDHαc2pro-NRSE-R | GGTAGGAGGGCTCAGCAAG |
| hPCDHγc3-NRSE-F† | TGAGCCGAAATGAATACT |
| hPCDHγc3-NRSE-R | GGCGCGCTCCAACACCAG |
| hPCDHγc4-NRSE-F† | GCGGCAGCAGCTGGACTT |
| hPCDHγc4-NRSE-R | CATCTGCATCCTGAGCCT |
| hPCDHγc5-NRSE-F† | ATCAGCAGCATCTGGGGCAC |
| hPCDHγc5-NRSE-R | CACAGTATTGGTGCCCAC |
| hHS7#1-NRSE-F† | GCTCTGAGGGCAACTAAAG |
| hHS7#1-NRSE-R | AAACCTCTGATTCGGCAC |
| hHS7#2-NRSE-F† | CATTTCTCTCTTTTGTCTCAG |
| hHS7#2-NRSE-R | CAGAATCTGCCTGTTTACC |
| hSE#1-NRSE-F† | TTCTAGGATGTGGGTGG |
| hSE#1-NRSE-R | GCTGACATACCTGAGTCCT |
| hSE#2-NRSE-F† | AGGCACTATCCCAAACAAGG |
| hSE#2-NRSE-R | TCTGCCCTCTAGCCTATTGT |
| hSE#3-NRSE-F† | ATTTCCTGACAGGTAGAGGG |
| hSE#3-NRSE-R | GATGTCATCACGTTAGAGAG |
| hSE#4-NRSE-F† | CTAAAATAAAAGGGATCCAT |
| hSE#4-NRSE-R | ATTTTATTTCTAGCAGTGTGC |
| hSE#5-NRSE-F† | CTCCAATAGTAAGTCTTCTGT |
| hSE#5-NRSE-R | GGGGCAAAACTTTAAAACT |
| hSE#6-NRSE-F† | TTCCCAGTGCTCAGCGT |
| hSE#6-NRSE-R | CTTATATCCCCAGCAGTTAATG |
| hSE#7-NRSE-F† | CTCCCCTTGGGAAACAG |
| hSE#7-NRSE-R | CATATTCAACCCCCCTGG |
| hSE#8-NRSE-F† | CCCCAGTGCCTCCTTGTGC |

| | |
|---|---|
| hSE#9-NRSE-F† | ATCAACTTTTGTTCATACACAC |
| hSE#9-NRSE-R | AAGCAGATATTTGGAACAAAG |
| hSE#10-NRSE-F† | GTGAAATCCCAAAAAGTCCATAG |
| hSE#11-NRSE-F† | CGGTTTCAAAAATAACAT |
| hSE#11-NRSE-R | AATCAAATGCCCCAAG |
| hSE#12-NRSE-F† | TCCTGCGTCACAAATAC |
| hSE#12-NRSE-R | CATAAAACAGACTCCCTATCT |
| hSE#13-NRSE-F† | CAGGCAGCTCCACCTAACCT |
| hSE#14-NRSE-F† | ACTGCTGTTTTTTTCCCTTCT |
| hSE#14-NRSE-R | CCCATCTCCACCCCTTC |
| hHS5-1- NRSE-F† | GCAGCGAGTCATGGGACC |
| hHS5-1- NRSE-R | TAAAGGAAATCTCTCTGGCC |
| Celsr3-I-NRSE-F† | TGTCTTCCAGGGGCCTCG |
| Celsr3-I-NRSE-R | AACCTGATGCAGGAGCTGTC |
| Celsr3-II-NRSE-F† | CTGCATCAGGTTCAGCACC |
| Celsr3-II-NRSE-R | AAGAGACCCCGGGAGCG |
| Celsr3-tandem-NRSE-F† | CTGGATTCAGCACCACGCAC |
| Celsr3-tandem-NRSE-R | GCTCGGGAGCTGTCCGAG |
| MAPK8IP-NRSE-F† | AACCAAGCCCAGGGCCGC |
| MAPK8IP-NRSE-R | CCCCACCCCGCCAGCCCC |
| synapsin1-NRSE-F† | GATGCGGCGAGGCGCGTG |
| synapsin1-NRSE-R | CGGTGGCGCGCGCCGCCA |
| hPCDHγa6-tandem-NRSE-F† | GCGCCCTGCTGGACAGAG |
| hPCDHγa6-tandem-NRSE-R | GCGGAGAGAGGGGGCTGG |
| hPCDHγa7-tandem-NRSE-F† | GGGCCCTGCTGGACAGAG |
| hPCDHγa7-tandem-NRSE-R | GCTGACAGAGGAGGCTGAC |
| Primers used to generate plasmids for shRNA knockdown | |
| REST-shRNA1-F | CCGGGCAAACACCTCAATCGCCATTCTCGAGAATGGCG ATTGAGGTGTTTGCTTTTTG |
| REST-shRNA1-R | AATTCAAAAAGCAAACACCTCAATCGCCATTCTCGAGA ATGGCGATTGAGGTGTTTGC |
| REST-shRNA2-F | CCGGCATGCAAGACAGGTTCACAATCTCGAGATTGTGA ACCTGTCTTGCATGTTTTTG |
| REST-shRNA2-R | AATTCAAAAACATGCAAGACAGGTTCACAATCTCGAGA TTGTGAACCTGTCTTGCATG |
| REST-shRNA3-F | CCGGGGAGCAAGTCCTTATTGAAGTCTCGAGACTTCAA TAAGGACTTGCTCCTTTTTG |
| REST-shRNA3-R | AATTCAAAAAGGAGCAAGTCCTTATTGAAGTCTCGAGA CTTCAATAAGGACTTGCTCC |
| GFP-shRNA-F | CCGGGTCGAGCTGGACGGCGACGTACTCGAGTACGTCG CCGTCCAGCTCGACTTTTTG |
| GFP-shRNA-R | AATTCAAAAAGTCGAGCTGGACGGCGACGTACTCGAGT ACGTCGCCGTCCAGCTCGAC |
| Primers used for construction of sgRNA expressing plasmids and for genotyping | |

| | |
|---|---|
| hHS51-NRSE-sgRNA1-F | ACCGACAGCGACACCGCCCAGTT |
| hHS51-NRSE-sgRNA1-R | AAACAACTGGGCGGTGTCGCTGT |
| hHS51-NRSE-sgRNA2-F | ACCGTGGCCATGGTGCTGAACTC |
| hHS51-NRSE-sgRNA2-R | AAACGAGTTCAGCACCATGGCCA |
| hHS51-NRSE-PCR-F | CTTGGAACCAGTTGGGATTG |
| hHS51-NRSE-PCR-R | TTATCAATAGCATTTTCCTCATCTG |
| mHS51-NRSE-sgRNA1-F | TAATACGACTCACTATAGGGCGAGTCATGGGACTCAACT |
| mHS51-NRSE-sgRNA2-F | TAATACGACTCACTATAGGATCTGGGGTGCTGAATCCT |
| Mouse-sgRNA-R | AAAAGCACCGACTCGGTGCC |
| mHS51-NRSE-PCR-F | GTTCCTTTGTCAGGTGAAAATCT |
| mHS51-NRSE-PCR-R | GGGTGGTAGTGAGGGATTATTCTAG |
| Primers used for QHR-4C | |
| 4C-adaptorF | GACGTGTGCTCTTCCGATCTGNNNNNN |
| 4C-adaptorR‡ | CAGATCGGAAGAGCACACGTC |
| 4C-hHS51-bioprimer† | TGCTTTCTCATTCCCCGTTG |
| 4C-hHS51-DpnII-F1 | AATGATACGGCGACCACCGAGATCTACACTCTTTCCCTACACGACGCTCTTCCGATCTCAGTTTTGGCGGCGACAAATTCG |
| 4C-hHS51-DpnII-F2 | AATGATACGGCGACCACCGAGATCTACACTCTTTCCCTACACGACGCTCTTCCGATCTCACGTTTTGGCGGCGACAAATTCG |
| 4C-hα12-bioprimer† | CCGCACCCACATTCCAATCA |
| 4C-hα12-DpnII-F1 | AATGATACGGCGACCACCGAGATCTACACTCTTTCCCTACACGACGCTCTTCCGATCTTCCAATCATTCACGGAATAGGATC |
| 4C-hα12-DpnII-F2 | AATGATACGGCGACCACCGAGATCTACACTCTTTCCCTACACGACGCTCTTCCGATCTGTCCAATCATTCACGGAATAGGATC |
| 4C-mHS51-bioprimer† | GCTTTGTTACTCTAGGAACAG |
| 4C-mHS51-DpnII-F1 | AATGATACGGCGACCACCGAGATCTACACTCTTTCCCTACACGACGCTCTTCCGATCTGGAGGTTAAAGCAAAGACTAAGATC |
| 4C-mHS51-DpnII-F2 | AATGATACGGCGACCACCGAGATCTACACTCTTTCCCTACACGACGCTCTTCCGATCTGAGGAGGTTAAAGCAAAGACTAAGATC |
| 4C- mα9-bioprimer† | GATATAGTTCGCTGTTTCTCAGGG |
| 4C-mα9-DpnII-F1 | AATGATACGGCGACCACCGAGATCTACACTCTTTCCCTACACGACGCTCTTCCGATCTTAGGAAGTAGCTACGTTCGGAG |
| P7-index-R1 | CAAGCAGAAGACGGCATACGAGATCTGCTAGTGACTGGAGTTCAGACGTGTGCTCTTCCGATCT |
| P7-index-R2 | CAAGCAGAAGACGGCATACGAGATGTCTAGGTGACTGGAGTTCAGACGTGTGCTCTTCCGATCT |

| | |
|---|---|
| P7-index-R3 | CAAGCAGAAGACGGCATACGAGATTCGTACGTGACTGG AGTTCAGACGTGTGCTCTTCCGATCT |
| P7-index-R4 | CAAGCAGAAGACGGCATACGAGATAGTACGGTGACTGG AGTTCAGACGTGTGCTCTTCCGATCT |
| P7-index-R5 | CAAGCAGAAGACGGCATACGAGATGACCTAGTGACTGG AGTTCAGACGTGTGCTCTTCCGATCT |
| P7-index-R6 | CAAGCAGAAGACGGCATACGAGATGAGTTCGTGACTGG AGTTCAGACGTGTGCTCTTCCGATCT |
| P7-index-R7 | CAAGCAGAAGACGGCATACGAGATGAAGCTGTGACTGG AGTTCAGACGTGTGCTCTTCCGATCT |
| P7-index-R8 | CAAGCAGAAGACGGCATACGAGATCTTGCAGTGACTGG AGTTCAGACGTGTGCTCTTCCGATCT |

† 5' biotin labeled

‡ phosphorylated
